# Differential contributions of β-tubulin isotypes to acentrosomal oocyte meiosis in *C. elegans*

**DOI:** 10.64898/2025.12.03.692175

**Authors:** Emmanuel T. Nsamba, Anne M. Villeneuve

## Abstract

Oocyte meiotic spindles must achieve bipolarity and segregate chromosomes in the absence of centrosomes. Here we use high-resolution immunofluorescence microscopy and live imaging to investigate the differential contributions of β-tubulin isotypes (TBB-1 and TBB-2) to assembly and function of acentrosomal spindles in *Caenorhabditis elegans* oocytes. By combining strains with altered β-tubulin isotype composition with mutations affecting microtubule-crosslinking motor KLP-18 and/or mutations affecting katanin-mediated microtubule severing, we show that TBB-1 and TBB-2 make distinct contributions to promoting spindle bipolarity. Further, by measuring multiple spindle features in wild-type and β-tubulin isotype substitution strains, we reveal contributions of isotype composition to spindle morphology, kinetics of anaphase chromosome separation and maintenance of spindle structural integrity under stress. Together, our data support a model in which β-tubulin isotype composition helps to maintain a balance between microtubule crosslinking and severing activities during oocyte meiosis. We further propose that this balance is crucial for establishing spindle bipolarity, maintaining spindle structures, and modulating the dynamics of chromosome separation.

## Introduction

Faithful chromosome segregation during meiosis requires the assembly of bipolar spindles, which physically separate chromosomes through two rounds of divisions to generate haploid gametes. In most metazoan cell types, centriole-containing centrosomes serve as dominant microtubule organizing centers that nucleate and organize spindle microtubules. In contrast, oocytes in most animals, including humans, *Drosophila*, and *C. elegans*, assemble functional spindles through acentrosomal mechanisms – a conserved but mechanistically distinct mode of spindle formation (Mogessie et al. 2018; Mullen et al. 2019). Failure to segregate chromosomes correctly during oocyte meiosis has a profound impact on human health, as it underlies a major fraction of pregnancy loss, infertility, and developmental disorders.

*Caenorhabditis elegans* has emerged as a powerful model system for dissecting acentrosomal mechanisms of spindle assembly, as meiotic divisions can be synchronized and visualized in intact organisms (Mullen et al. 2019; Wolff et al., 2022). Pioneering studies established that acentrosomal spindle formation proceeds through a sequence of coordinated steps (Wolff et al., 2016; Chuang et al. 2020) mediated by the dynamic interplay between microtubules and an ensemble of microtubule-interacting factors, including molecular motors and proteins that regulate microtubule dynamics (Fig 1A). Central among these factors is the plus-end-directed homodimeric kinesin-12 motor KLP-18, which crosslinks microtubules to establish and maintain spindle bipolarity (Segbert et al. 2003); KLP-18 depletion results in monopolar spindles and embryonic lethality (Wignall and Villeneuve, 2009). Also critical is the microtubule-severing enzyme katanin, comprising the MEI-1 catalytic subunit and MEI-2 regulatory subunit in *C. elegans* (Mains et al. 1990; Clark-Maguire and Mains 1994). Katanin functions as a central orchestrator of multiple aspects of acentrosomal spindle assembly in *C. elegans*, contributing to spindle pole formation and organization, chromosome alignment, and regulation of microtubule length and dynamics (Srayko et al. 2000; 2006; Connolly et al. 2014; McNally et al. 2014; Lantzsch et al. 2021).

**Figure 1.**
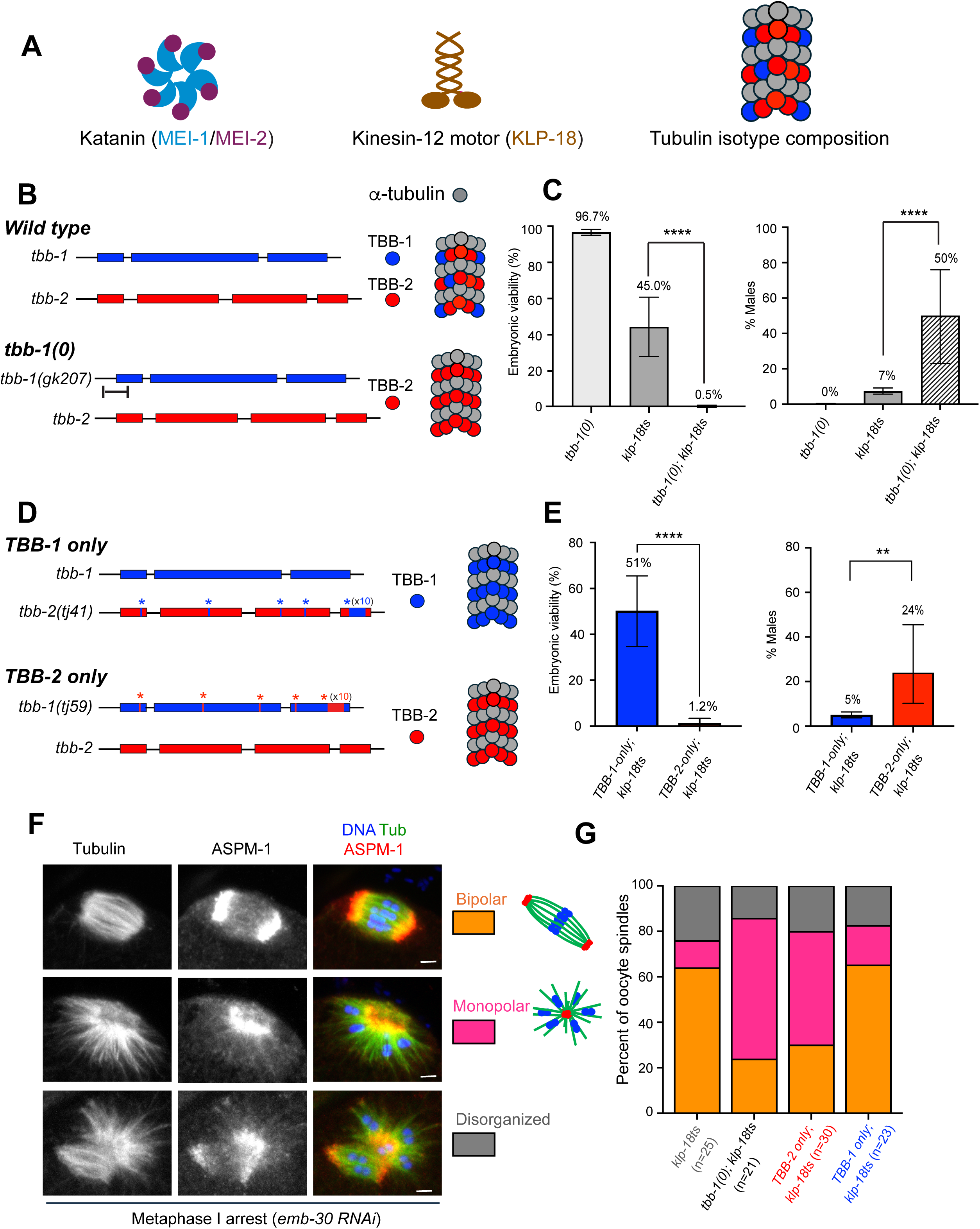
Differential requirements for β-tubulin isotypes TBB-1 and TBB-2 in promoting bipolarity of oocyte meiotic spindles. **(A)** Schematic illustrating key spindle assembly factors featured in this work: heterodimeric microtubule severing enzyme katanin composed of enzymatic subunit MEI-1 and regulatory subunit MEI-2; plus-end directed kinesin-12 motor KLP-18; and microtubules containing two distinct β-tubulin isotypes,TBB-1 (blue) and TBB-2 (red). **(B)** Diagrams depicting the *tbb-1* and *tbb-*2 gene structures (for the wild-type and *tbb-1* null genotypes), the β-tubulin isotypes produced for each genotype, and the composition of resulting microtubules. **(C)** Graphs reporting frequencies of viable embryos (left) and of male progeny (indicative of X-chromosome mis-segregation, right) produced by hermaphrodites of the indicated genotypes at 20°C (semi-permissive temperature for the *klp-18ts* mutation); these data demonstrate a synthetic genetic interaction between *tbb-1(0)* and *klp-18ts*. For embryo viability graph, error bars indicate standard deviation; for male frequency graph, error bars indicate 95% confidence interval. Data were analyzed using two-tailed Mann-Whitney test and Fisher exact test, respectively; **p < 0.01; ****p < 0.0001. **(D)** Gene structure and microtubule composition diagrams for the β-tubulin isotype substitution strains, which express normal levels of β-tubulin. **(E)** Graphs revealing isotype-specific effects on embryo viability and male frequency in the *klp-18ts* mutant background; error bars and p values as in C. **(F)** Immunofluorescence (IF) images of *klp-18ts* spindles arrested at metaphase I (following *emb-30 RNAi*), showing representative examples of the different spindle morphologies observed; scale bars, 2 µm. **(G)** Quantification of spindle morphologies observed in F; n = number of spindles. p values were calculated using Fisher exact test with two categories: “bipolar” vs. “monopolar + disorganized”. Significant differences were observed for: *tbb-1(0); klp-18ts* vs *klp-18ts,* p=0.009; *TBB-2-only; klp-18ts* vs *klp-18ts,* p=0.015; *TBB-2-only; klp-18ts* vs *TBB-1-only; klp-18ts,* p=0.014; *tbb-1(0); klp-18ts* vs *TBB-1-only; klp-18ts,* p=0.008.

The coordinated activities of KLP-18, katanin, and other microtubule-interacting proteins during acentrosomal spindle assembly necessitate their productive engagement with microtubules that are compositionally heterogeneous, containing multiple variants, or isotypes, of α- and β-tubulin subunits. Accumulating evidence across diverse systems indicates that tubulin isotypes confer distinct functional properties to microtubule-dependent processes (Nsamba and Gupta 2022). During mitosis in budding yeast, for example, α-tubulin isotypes exhibit specialization for distinct spindle positioning mechanisms (Nsamba et al. 2021) and contribute divergent properties that balance anaphase spindle dynamics (Nsamba et al. 2025). Similarly, studies in *C. elegans*, *Drosophila*, and mice have revealed isotype-specific contributions to neurite growth (Lockhead et al. 2016; Zheng et al. 2017; Baran et al. 2010), cilia function (Silva et al. 2017; Hurd et al. 2010.), temperature adaptation (Myachina et al. 2017), spermatogenesis (Hutchens et al. 1997; Hoyle and Raff 1990), and neuronal development (Bittermann et al. 2019; Buscaglia et al. 2020) and axonal regeneration (Latremoliere et al. 2018)

The *C. elegans* system, which encodes two germline-expressed β-tubulin isotypes, TBB-1 and TBB-2 (Nishida et al. 2021) provides an outstanding opportunity to investigate the functional significance of tubulin isotype diversity to the formation and function of acentrosomal meiotic spindles. β-tubulin isotype substitution strains have been created that express wild-type levels of tubulin but produce only one of the two isotypes, permitting direct investigation of potential functional differences (Honda et al. 2017). At a superficial level, these two β-tubulin isotypes appear at least partially functionally redundant, as embryos from the tubulin isotype substitution strains are fully viable. Despite this apparent redundancy, genetic analyses involving mutations that either partially reduce or cause ectopic persistence of katanin activity have provided evidence that the microtubules containing the TBB-2 β-tubulin isotype are preferred substrates for katanin-mediated severing *in vivo* (Lu et al. 2004). However, the extent of functional specialization of the TBB-1 and TBB-2 isotypes and their specific contributions to meiotic spindle assembly and function have remained largely unexplored.

Here, we employ high-resolution immunofluorescence microscopy and live imaging in *Caenorhabditis elegans* strains with altered β-tubulin isotype composition to investigate how TBB-1 and TBB-2 differentially contribute to acentrosomal spindle assembly and chromosome segregation dynamics. Through a sensitized genetic approach, we provide evidence that TBB-1 and TBB-2 make distinct contributions to a system that counterbalances microtubule crosslinking and microtubule severing activities to enable stable formation of bipolar meiotic spindles. Further, by comparing multiple features of meiotic spindles from wild type and β-tubulin isotype substitution strains, we reveal hidden contributions of isotype composition to spindle length, katanin localization, and kinetics of anaphase chromosome separation. Overall, these findings reveal how tubulin isotype diversity enables fine-tuned temporal and spatial control of acentrosomal spindle function.

## Results

### Absence of the TBB-1 isotype exacerbates oocyte meiotic defects caused by *klp-18ts*

To test whether a sensitized genetic background might be useful for revealing potential differential contributions of β-tubulin isotypes to assembly and/or function of meiotic spindles, we employed *klp-18(or447ts)*, a temperature-sensitive missense mutation causing partial loss-of-function of KLP-18 (Connolly et al. 2014), a plus-end-directed kinesin-12 microtubule crosslinking motor required for bipolarity of oocyte meiotic spindles (Segbert et al. 2003). Through construction of double mutants, we discovered a synthetic genetic interaction between a *tbb-1(0)* null mutation and *klp-18(or447ts)* at the semi-permissive temperature of 20°C (Table S1 and Figure 1B-C). Whereas *tbb-1(0)* and *klp-18ts* single mutants exhibited 97% and 45% embryonic viability, respectively, nearly all embryos produced by *tbb-1(0); klp-18ts* double mutant animals were inviable (0.5% viability; p < 0.0001 compared to either single mutant). Further, 50% of the rare surviving progeny were males (compared to 0.1% and 7% for the respective single mutants; p < 0.0001), a phenotype indicative of meiotic missegregation of sex chromosomes.

To further investigate this synthetic interaction, we introduced the *klp-18ts* mutation into β-tubulin substitution strains generated by CRISPR editing to enable expression of single isotypes at normal protein levels (Honda et al. 2017). “TBB-1-only” strains express TBB-1 from both the *tbb-1* and *tbb-2* loci, while “TBB-2-only” strains express TBB-2 from both loci, thus maintaining normal total β-tubulin levels while altering isotype composition of the microtubules (Honda et al. 2017; Figure 1D). We found that *TBB-1-only; klp-18ts* double mutant worms exhibited levels of embryonic viability and incidence of male progeny similar to those observed for the *klp-18ts* single mutant (Table S1; Figure 1E). In contrast, *TBB-2-only; klp-18ts* double mutant worms exhibited severe synthetic defects comparable to the original *tbb-1; klp-18ts* synthetic interaction: 1.2% embryo viability and 24% male progeny (Table S1; Figure 1E). Together, these data suggest that the presence of TBB-1 in microtubules is required for oocyte meiotic spindle function when KLP-18 activity is partially impaired.

To understand the cellular basis for these genetic interactions, we used immunofluorescence microscopy to directly assess the morphology of metaphase I-arrested oocyte meiotic spindles (Furuta et al. 2000), using antibodies against ASPM-1 as a spindle pole marker (Wignall and and Villeneuve 2009) and anti-α-tubulin antibodies to visualize microtubules (Figure 1F-G, Supplemental Figure 1). Whereas the majority of spindles in the *klp-18ts* single mutant and the *TBB-1-only; klp-18ts* double mutant exhibited normal bipolar organization, double mutants lacking the TBB-1 isotype (both *tbb-1(0); klp-18ts* and *TBB-2 only; klp-18ts*) exhibited elevated frequencies of monopolar spindles. Analysis of meiotic spindles in unsynchronized embryos similarly revealed an elevated incidence of monopolar spindles in the *TBB-2-only; klp-18ts* double mutant relative to the *klp-18ts* single mutant (Supplemental Figure 2A-D). We conclude that microtubules containing only the TBB-2 β-tubulin isoform are deficient in promoting bipolarity of acentrosomal oocyte meiotic spindles under conditions where KLP-18-dependent microtubule crosslinking activity is compromised.

In principle, worsening of the *klp-18* mutant phenotype could potentially be caused by reduced recruitment of the mutant KLP-18 protein. However, immunostaining experiments did not detect differences in KLP-18 localization between TBB-1-only versus TBB-2-only spindles (Supplemental Figure 3A-B), suggesting that impaired KLP-18 recruitment is unlikely to be responsible for the observed differences in genetic sensitivity to the *klp-18ts* mutation.

### Mutations causing either reduced sensitivity to or reduced activity of katanin suppress the synthetic meiotic defects of *TBB-2 only; klp-18ts*

Based on the prior work of Lu et al., 2004, we hypothesized that the synthetic meiotic defects observed in the *TBB-2-only; klp-18ts* double mutant might result from elevated sensitivity of TBB-2-only microtubules to the microtubule severing enzyme katanin. We tested this hypothesis in two ways.

First, we used a missense mutation, *tbb-2(E439K)*, that was initially isolated based on suppression of lethality caused by ectopic persistence of katanin activity in cleavage-stage embryos and is inferred to confer partial resistance of microtubules to katanin-mediated severing (Lu et al. 2004). We found that introduction of the *tbb-2(E439K)* missense mutation into the *TBB-2-only; klp-18ts* background (Figure 2A) was indeed sufficient to suppress the embryonic lethality, sex chromosome missegregation, and meiotic spindle defects observed in the *TBB-2 only; klp-18ts* double mutant (Table S2 and Figure 2B-E).

**Figure 2.**
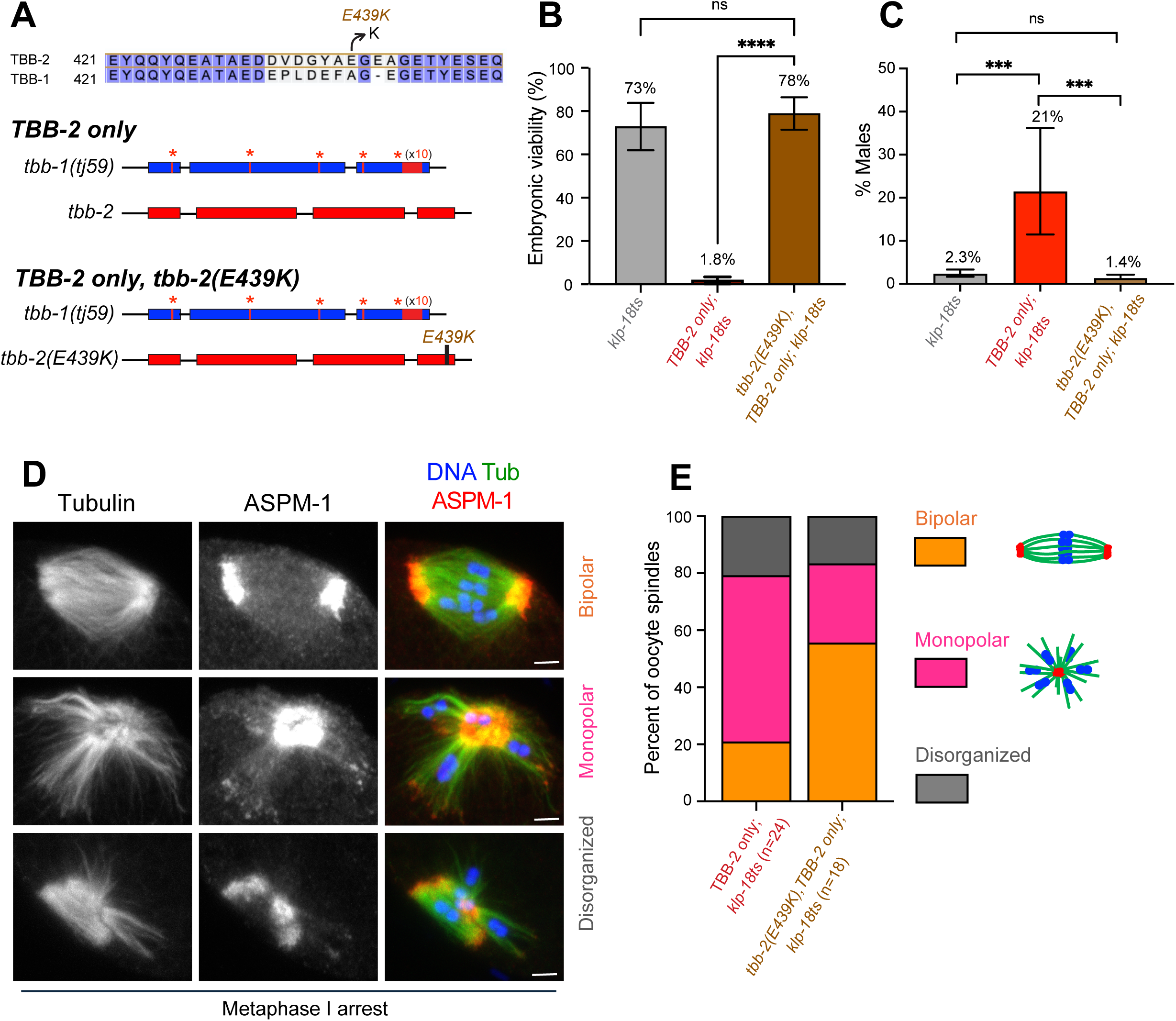
The *tbb-2(E439K)* mutation, inferred to confer partial resistance to katanin-mediated severing, suppresses synthetic meiotic defects of *TBB-2 only; klp-18ts*. **(A)** Top, protein sequence alignment showing E439K position in TBB-2 carboxy-terminal tail. Lavender shading shows identical residues. Bottom, gene structure diagrams showing the location of the *tbb-2(E439K)* missense mutation in the *tbb-2* locus. **(B, C)** Quantification of embryonic viability and male frequency at 20°C, as in Figure 1C. **(D)** Representative IF images of metaphase I arrested oocyte spindles in the *tbb-2(E439K)* mutant background; scale bars, 2 µm. **(E)** Quantification of spindle phenotypes observed in D; n = number of spindles. Analysis of spindle bipolarity as in Figure 1G, p=0.048.

Second, we sought to test whether reducing katanin activity itself might similarly suppress the *TBB-2 only; klp-18ts* meiotic defects. For this purpose, we used mei-2(A237T), a partial loss-of-function allele affecting katanin accessory subunit MEI-2 (Srayko et al. 2000) that was previously shown to reduce katanin severing activity *in vitro* without affecting embryonic viability (McNally et al. 2014) (Figure 3A). Introduction of the *mei-2(A237T)* missense mutation into the *TBB-2 only; klp-18ts* mutant background provided robust suppression comparable to that observed with *tbb-2(E439K)* (Table S3 and Figure 3B-D). Critically, *mei-2(A237T)* substantially restored bipolar spindle formation to levels similar to those observed for the *klp-18ts* single mutant.

**Figure 3.**
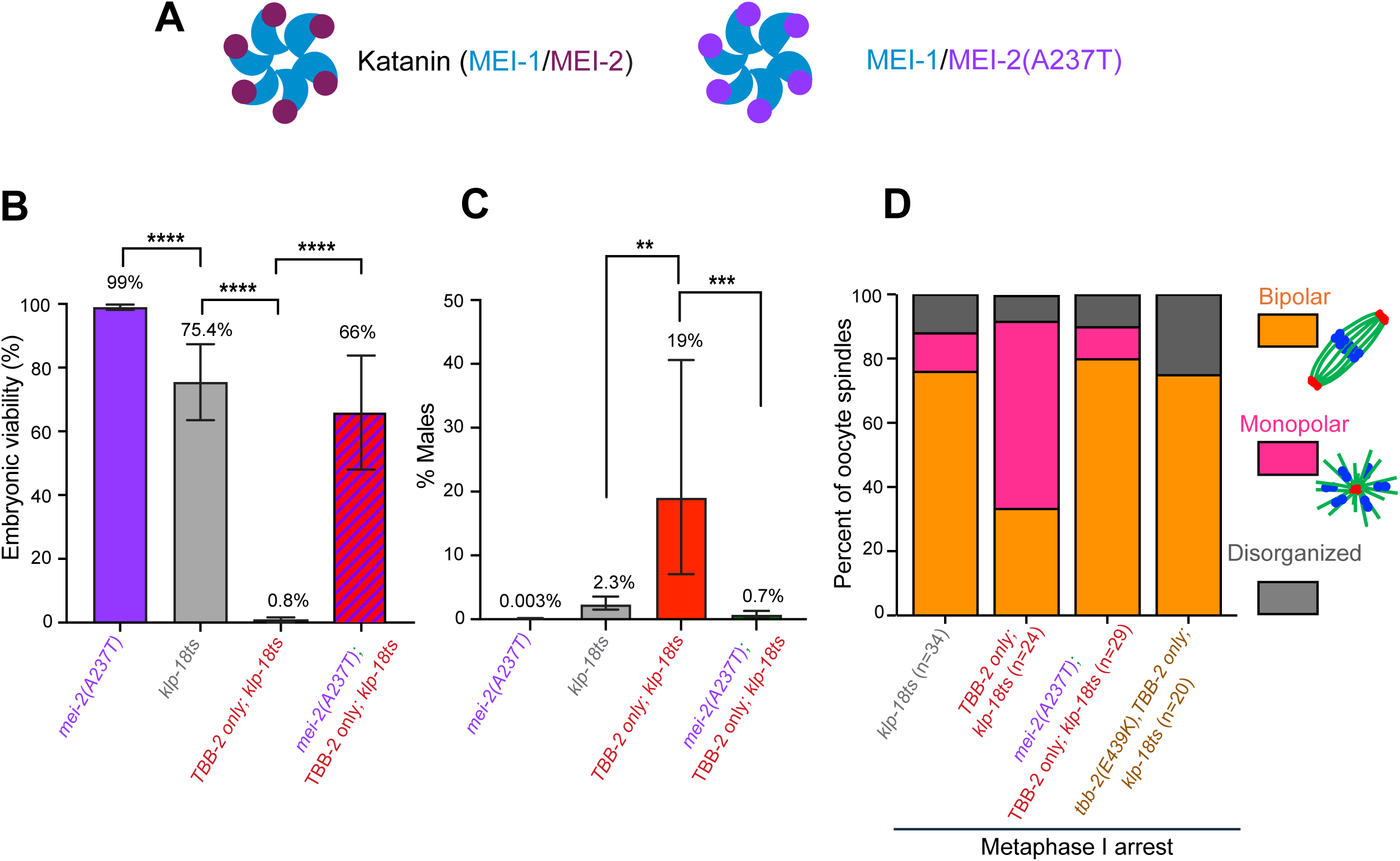
Reduction of katanin activity suppresses synthetic meiotic defects of *TBB-2 only; klp-18ts.* **(A)** Schematic of katanin in wild type and in the *mei-2(A237T)* partial loss-of-function mutant (ref). **(B, C)** Quantification of embryonic viability and male frequency at 20°C, as in Figure 1C. *mei-2(A237T)* restores embryonic survivability and suppresses chromosome missegregation of *TBB-2 only; klp-18ts*. **(D**) Quantifications of morphologies observed in IF images of metaphase I arrested oocyte spindles; n = number of spindles. Analysis of spindle bipolarity as in Figure 1G. Significant differences were observed for: *TBB-2-only; klp-18ts* vs *klp-18ts*, p =0.001; *TBB-2-only; klp-18ts* vs *tbb-2(E439K),TBB-2-only; klp-18ts,* p=0.003; *TBB-2-only;klp-18ts* vs *mei-2(A437T); TBB-2-only; klp-18ts*, p=0.002.

Together, our data showing suppression both by reducing sensitivity of microtubule to katanin and by reducing katanin activity support our hypothesis that the enhanced meiotic defects observed in *TBB-2-only; klp-18ts* oocytes reflect hypersensitivity of TBB-2-only microtubules to katanin-mediated severing. Our data suggest that β-tubulin isotype composition promotes bipolarity of *C. elegans* oocyte meiotic spindles in part by modulating susceptibility of spindle microtubules to katanin-mediated severing, thereby helping to maintain a crucial balance between microtubule stabilizing (e.g. crosslinking) and destabilizing (e.g. severing) activities.

### β-tubulin isotype composition differentially affects association of katanin with meiotic spindles

To further investigate the relationship between β-tubulin isotype composition and katanin, we examined the localization of MEI-2 (katanin regulatory subunit) and MEI-1 (katanin enzymatic subunit) in metaphase I-arrested oocyte spindles from wild-type, *TBB-1-only*, and *TBB-2-only* strains (in a wild-*type klp-18(+)* background (Figure 4, Supplemental Fig S4). Previous studies showed that katanin localizes to *C. elegans* oocyte spindle poles, where it promotes pole organization, and also localizes with chromosomes in the middle of the spindle, although the functional significance of chromosome-localized katanin is unclear (Srayko et al. 2000; McNally et al. 2014). We similarly observed spindle pole localization of MEI-2 and MEI-1 in all genotypes, but we also noted that *TBB-1-only* oocyte spindles appeared to have reduced chromosome-localized MEI-2 and MEI-1 in their central spindles.

**Figure 4.**
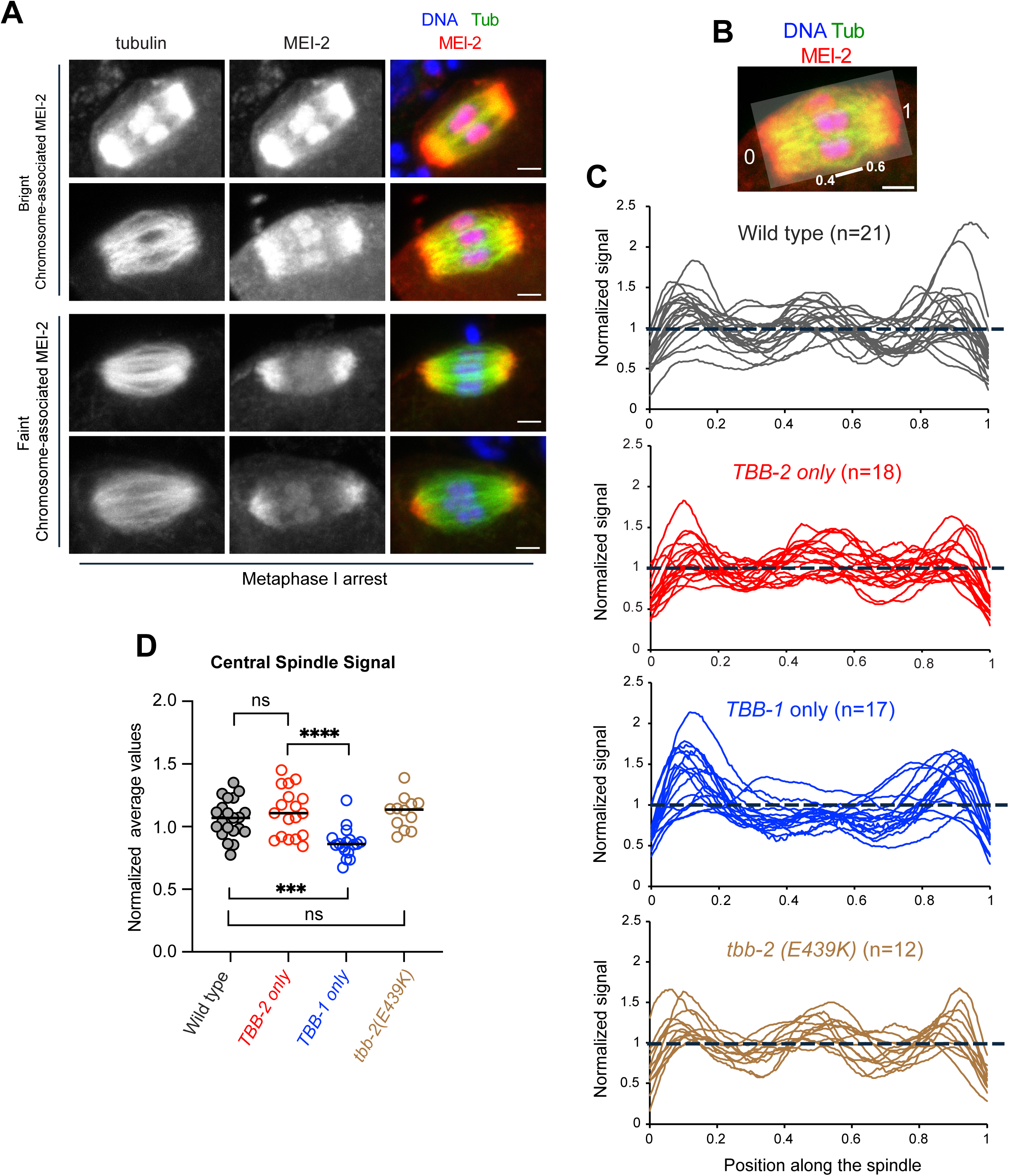
β-tubulin isotype composition differentially affects association of katanin with meiotic spindles. **(A)** Immunolocalization of MEI-2 at the poles and around the chromosomes in metaphase I-arrested oocyte spindles. Images illustrate the range of patterns observed; scale bar = 2.0 μm. **(B)** Depiction of scheme for quantifying the relative MEI-2 distribution along the spindle axis (from 0-1), using a “line” tracing method wherein the width of the line corresponds to the full width of the spindle (see Materials and Methods). The region from 0.4 – 0.6 represents the central portion of the spindle in the vicinity of chromosomes. **(C)** Pileups of individual line traces of normalized MEI-2 signal intensity from measurements conducted as in B (and Materials and Methods). **(D)** Quantification of the normalized average MEI-2 signal in the chromosome-occupied central spindle (positions 0.4–0.6); each data point represents an individual spindle. Horizontal lines indicate median. MEI-2 signal is significantly depleted from central spindles in *TBB-1 only*. Data were analyzed using two-tailed Mann-Whitney test; ns, not significant; ***p < 0.001; ****p < 0.0001.

We conducted line profile analyses to quantitatively evaluate the spatial distribution of MEI-2 and MEI-1 along the spindle long axis (Figure 4, Supplemental Fig S4; see Materials and Methods). In these analyses, signal intensities at a given position along the long axis of a spindle are normalized to the mean intensity for that spindle, enabling evaluation of the distribution of signal in the chromosome-occupied central portion of the spindle relative to the pole regions. Composite plots of line profiles confirmed the enrichment of MEI-2 and MEI-1 near spindle poles for all three genotypes (Figure 4C, Supplemental Fig S4C ). Moreover, our analysis allowed us to conclude that normalized MEI-2 and MEI-1 signals near chromosomes in the central region of *TBB-1-only* spindles were significantly reduced compared to both *TBB-2-only* and wild-type spindles(Figure 4D, Supplemental Fig S4D ). Interestingly, MEI-2 localization profiles in *tbb-2(E439K)* spindles were not distinguishable from wild-type controls, consistent with previous reports suggesting that the TBB-2(E439K) amino acid change impairs microtubule-katanin interactions without disrupting katanin recruitment (Lu et al. 2004).

These findings regarding katanin localization help to provide a potential mechanistic explanation for the observed differences in sensitivity of spindle structure to partial loss of KLP-18 function: By recruiting less katanin to the spindle center, TBB-1-containing microtubules may create regions of reduced severing activity, potentially leading to longer antiparallel overlaps that preserve spindle stability when crosslinking is compromised. Conversely, TBB-2-containing microtubules may recruit more katanin, creating higher local severing activity that becomes destructive when crosslinking is reduced – presumably by leading to shortened regions of antiparallel microtubule overlap.

### β-tubulin isotype composition differentially affects meiotic spindle organization

The spatial distribution of katanin predicted that TBB-1-only and TBB-2-only spindles might exhibit different structural properties. To test this possibility, we evaluated several aspects of spindle organization in high-resolution immunofluorescence images of α-tubulin and spindle pole marker ASPM-1 in metaphase I-arrested oocyte spindles in a wild-type [*klp-18*(+)] background. First, we found that TBB-1-only spindles were significantly longer than both wild-type and TBB-2-only spindles: 8.41 ± 0.56 μm versus 7.30 ± 0.53 μm (wild-type, p < 0.0001) and 7.20 ± 0.54 μm (TBB-2-only, p < 0.0001) (Figure 5A-B). Notably, mutants with partially reduced katanin activity [*gfp::mei-1; mei-2(A237T)]* or reduced katanin sensitivity [*tbb-2(E439K)]* similarly exhibited longer meiotic spindles, consistent with the possibility that reduced katanin activity/susceptibility may at least partially underlie the TBB-1-only phenotype.

**Figure 5.**
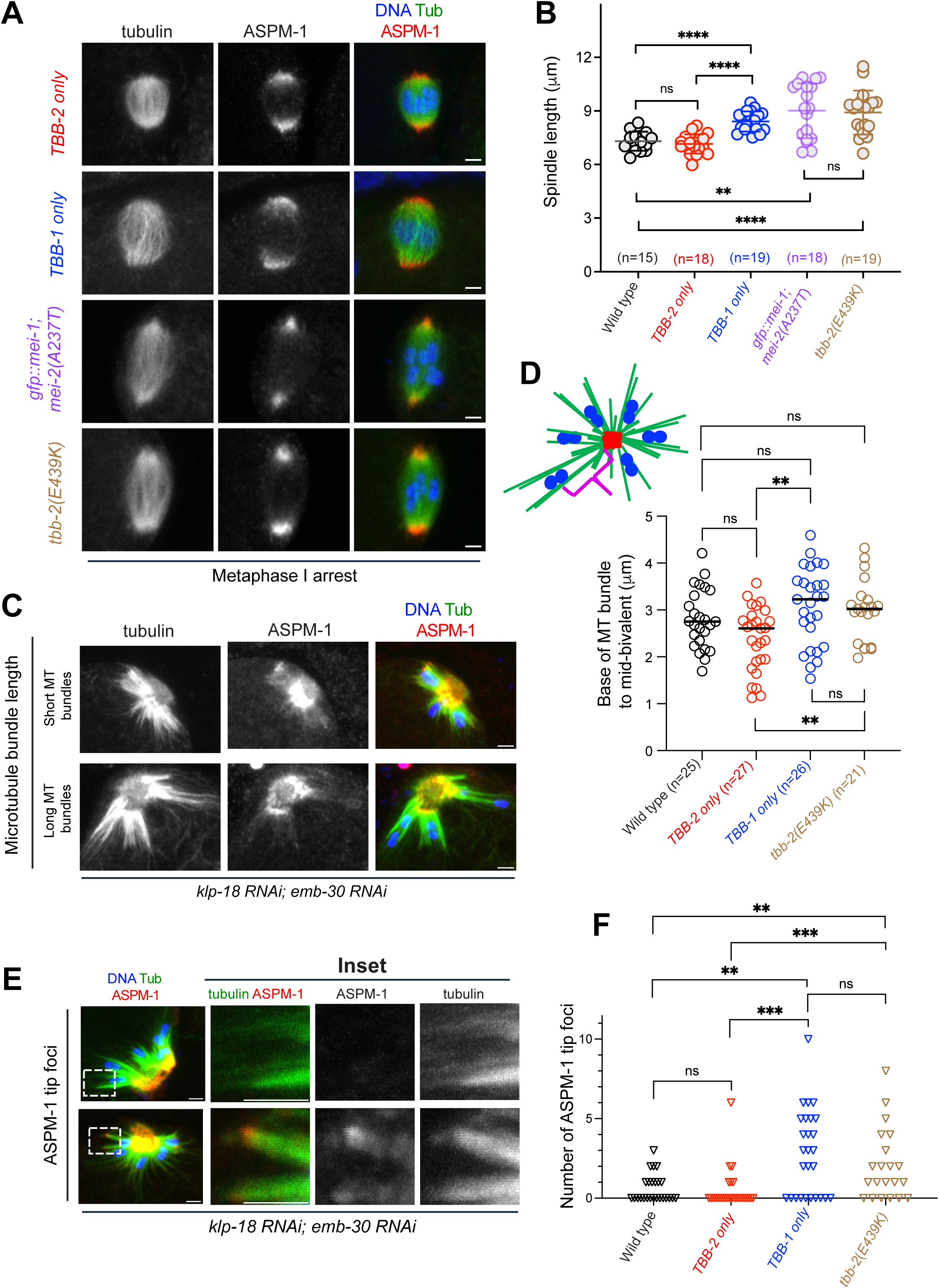
β-tubulin isotype composition differentially affects meiotic spindle morphology and properties of microtubules bundles. **(A)** Representative IF images of metaphase I arrested spindles illustrating length differences for the indicated genotypes; scale bar = 2.0 μm. **(B)** Quantification of spindle length measurements in A; n = number of spindles. Error bars indicate standard deviation. Data were analyzed using two-tailed Mann-Whitney test; ns, not significant; **p < 0.01; ****p < 0.0001. **(C)** IF images of metaphase I arrested monopolar spindles (*klp-18 RNAi*; *emb-30 RNAi*) showing bivalent positions that reflect microtubule bundle lengths. **(D)** Quantification of distances from images of monopolar spindles of the types depicted in C, measured from the base of a microtubule bundle to the bivalent midpoint, as shown in the schematic diagram. Each data point represents the mean of the individual base-to-mid-bivalent measurements from a given spindle. Horizontal lines indicate median. Data were analyzed using two-tailed Mann-Whitney test; ns, not significant; **p < 0.01. **(E)** IF images of monopolar spindles showing ASPM-1 tip foci (inset), which are indicative of antiparallel microtubule organization in bundles despite depletion of KLP-18. Scale bar = 2 μm (regular images); 10 μm (inset) **(F)** Quantification of the number of ASPM-1 tip foci observed at the periphery of each monopolar spindle. Each data point represents an individual monopolar spindle. Data were analyzed using two-tailed Mann-Whitney test; ns, not significant; **p < 0.01; ***p < 0.001.

We also evaluated spindle pole organization, assessing both pole width and total ASPM-1 signal associated with poles (Supplemental Figure S5). Although comparisons of wild-type, TBB-1-only and TBB-2 spindles did not reveal significant differences in total ASPM-1 signal, both TBB-1 only and TBB-2 only spindles had slightly narrower poles than wild-type. However, these modest differences contrasted sharply with pronounced defects in spindle pole organization detected in the *gfp::mei-1; mei-2(A237T)* and *tbb-2(E439K)* mutants, which displayed diminished ASPM-1 signal and narrower spindle pole structures.

### β-tubulin isotype composition differentially affects properties of microtubule bundles

We used monopolar spindles generated by depletion of KLP-18 by RNAi to seek additional evidence for differential microtubule properties affected by β-tubulin isotypes (Figure 5C-F). In this monopolar spindle configuration, lateral associations with microtubules align chromosome bivalents roughly parallel to groups of microtubule bundles that emanate from a region of high microtubule density in the area of the monopole (marked by high ASPM-1 signal). In this context, bivalents are transported away from the monopole towards the faint bundle tips via chromokinesin activity concentrated at the mid-bivalent (Wignall and and Villeneuve 2009); thus, we reasoned that the distance from the base of the microtubule bundles to the mid-bivalent could serve as a quantifiable surrogate measure reflecting relative length of microtubule bundles (see Material and Methods). TBB-1-only monopolar spindles did indeed exhibit significantly greater base-to-mid-bivalent distances compared to TBB-2-only monopolar spindles (3.47 ± 0.95 μm versus 2.78 ± 0.82 μm; p < 0.01) (Figure 5D), suggesting longer microtubule bundles and consistent with the longer spindle lengths measured for TBB-1-only bipolar spindles. Base-to-mid-bivalent distances in *tbb-2(E439K)* monopolar spindles were also significantly longer than in TBB-2-only (p = 0.0091), consistent with longer spindle lengths when microtubule susceptibility to katanin activity is reduced. Further, closer scrutiny of ASPM-1 localization in these monopolar spindles revealed that TBB-1-only monopolar spindles had a higher frequency of microtubule bundles with ASPM-1 foci at their peripheral tips (distal to the monopole), presumably indicative of antiparallel microtubule organization within such bundles (Figure 5E-F).

### Live imaging reveals hidden contributions of β-tubulin isotype composition to spindle morphology and chromosome separation dynamics

To investigate β-tubulin isotype contributions to the dynamics of meiotic spindle assembly and function, we used spinning disk confocal microscopy to conduct time-lapse *in utero* imaging of control, TBB-1-only, and TBB-2-only oocytes expressing GFP-tagged α-tubulin (GFP::TBA-2) and mCherry::H2B, to mark microtubules and chromosomes, respectively (Figure 6). These experiments build on pioneering live-imaging studies of *C. elegans* oocyte meiosis (Chuang et al. 2020) that defined landmark events and provide a framework for dissecting the roles of individual molecular components in both spindle formation and chromosome segregation dynamics. For our analyses of progression through meiosis I, we designated the appearance of microtubules surrounding the chromosomes (“Cage” stage) as the onset of meiotic spindle assembly (t=0), and we used the timing of Metaphase, Anaphase A onset, Anaphase B onset, and Maximum extent of chromosome segregation as landmarks to align and compare morphology and timing for control, TBB-1-only and TBB-2-only spindles (Figure 6A; Supplemental Figure 6; Materials and Methods). Several interesting observations emerged from these analyses:

**Figure 6.**
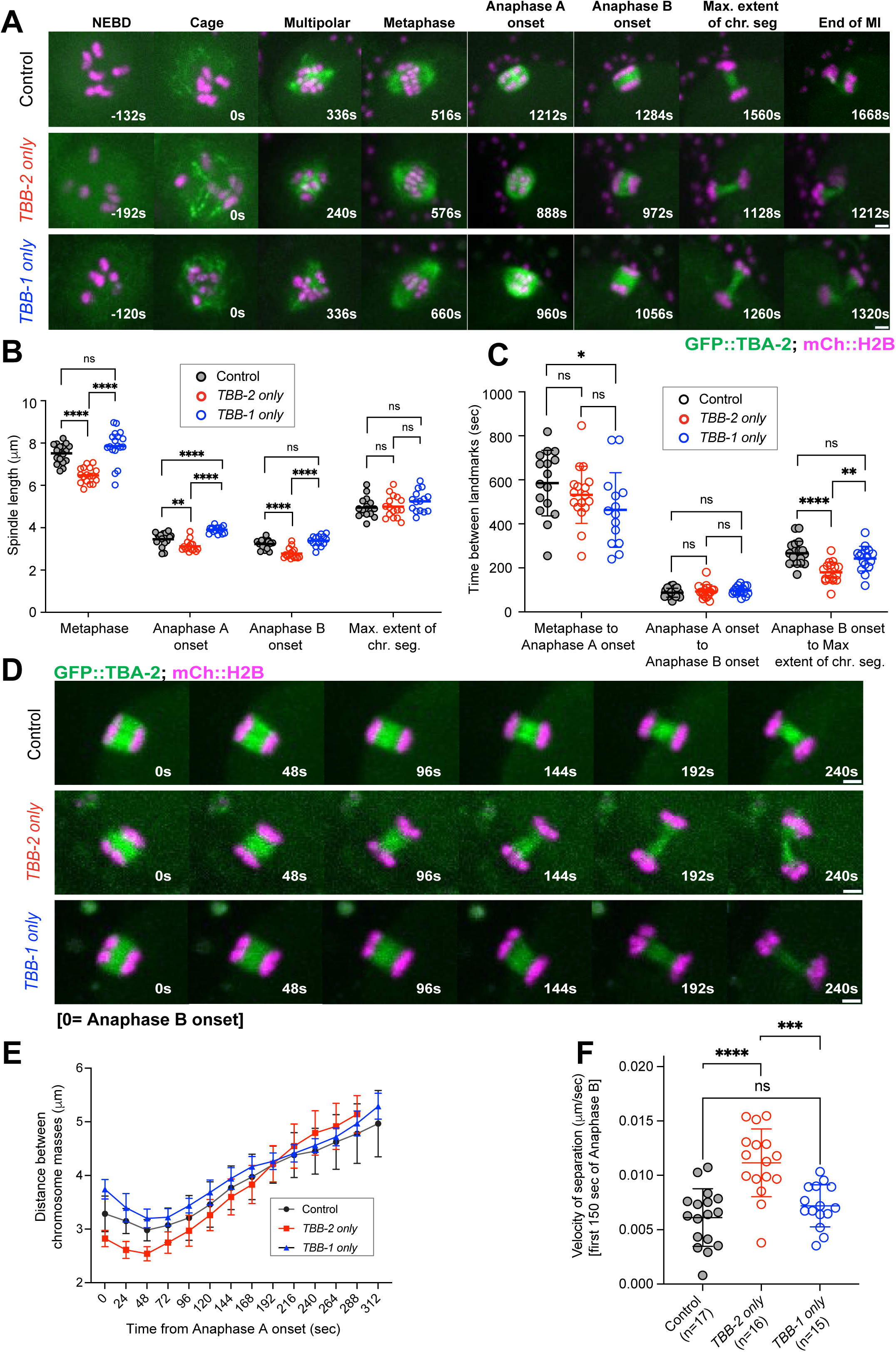
Live imaging reveals contributions of β-tubulin isotype composition to spindle morphology and chromosome separation dynamics. **(A)** Representative maximum intensity projection images from time-lapse movies of live oocytes expressing GFP::TBA-2 to mark microtubules and mCherry::H2B to mark chromosomes, showing meiosis I landmarks. For our analysis, t = 0 is the onset of meiotic spindle assembly as defined by the appearance of microtubules surrounding the chromosomes (“Cage” stage). Scale bar = 2.0 μm. **(B)** Quantification of spindle length at specified meiotic landmarks; error bars indicate standard deviation. **(C)** Timing of meiosis I spindle assembly measurements between specified meiotic landmarks; error bars indicate standard deviation. **(D)** Representative maximum intensity projection images from time-lapse movies of Anaphase B. Anaphase B onset (t = 0) is defined as the point when the spindle has achieved its shortest length and the separating chromosome masses have reached opposite poles of the spindle. **(E)** Quantification of chromosome separation distances (mean ± standard deviation) over time during anaphase I. **(F)** Quantification of anaphase B chromosome separation velocity during meiosis I for the first 150 seconds after anaphase B onset (see Materials and Methods) ; n = number of spindles analyzed. Error bars indicate standard deviation. For B, C and F, each data point represents a spindle, and data were analyzed using two-tailed Mann-Whitney test; ns, not significant; *p < 0.05; **p < 0.01; ***p < 0.001; ****p < 0.0001.

*TBB-2-only metaphase spindles are short but assemble with normal timing:* In these live imaging experiments, TBB-2-only metaphase spindles were significantly shorter than *both* control spindles (6.47 ± 0.36 μm versus 7.51 ± 0.45 μm; p < 0.0001) and TBB-1-only spindles (7.86 ± 0.77 μm; p < 0.0001) (Figure 6B). We note that this finding is only partially consistent with our measurements of spindle lengths in immunofluorescence imaging of metaphase I-arrested oocyte spindles (Figure 5), where we detected a length difference between TBB-2-only spindles and TBB-1-only spindles, but not between TBB-2-only spindles and those from wild-type controls; this apparent discrepancy is likely caused by the prolonged arrest, as discussed below. Despite differences in metaphase spindle lengths, we did not detect differences in the time to assemble bipolar metaphase spindles (Supplemental Figure 6A).

*Despite assembling shorter spindles, TBB-2-only oocytes achieved a normal extent of chromosome separation during anaphase:* The lengths of TBB-2-only spindles were significantly shorter than control and TBB-1-only spindles not only at metaphase, but also at both anaphase A onset and anaphase B onset. Nevertheless, TBB-2-only spindles ultimately achieved maximum chromosome separation distances comparable to those of the other genotypes (approximately 5.0 μm in all cases) (Figure 6B)

*TBB-2 only-microtubules promote accelerated spindle elongation and increased chromosome separation velocity during anaphase B:* TBB-2-only spindles reached maximum anaphase B chromosome separation significantly faster than control and TBB-1-only spindles: 180 ± 46 seconds from anaphase B onset versus 267 ± 54 seconds in control (p < 0.0001) and versus 242 ± 56 seconds in TBB-1-only oocytes (p = 0.0012) (Figure 6C). This reflects an elevated rate of chromosome separation during anaphase B in TBB-2-only oocytes, as revealed by monitoring of chromosome separation distances throughout anaphase B (Figure 6D) and visualized by plotting composite measurements of distance vs. time (Figure 6E) or by plotting average separation velocity measurements for individual spindles (Figure 6F and Supplemental Figure 6D).

### TBB-1-only spindles exhibit increased susceptibility to structural failure during prolonged arrest

We set out to investigate the basis of differences between our live imaging of oocyte meiosis and immunofluorescence imaging of metaphase-arrested spindles regarding effects of β-tubulin isotype composition on metaphase spindle length. To this end, we imaged GFP::TBA-2 and mCherry::H2B fluorescence in fixed metaphase I-arrested oocytes from the strains we had used for live imaging. This analysis largely recapitulated our previous finding that metaphase-arrested TBB-1-only spindles are significantly longer than both wild-type and TBB-2-only spindles (Supplemental Figure 7A-B), supporting the interpretation that prolonged arrest may exacerbate the effects of a modest functional difference between mixed-isotype and TBB-1-only microtubules. Notably, we also observed additional structural abnormalities (*e.g.* fractured or bent spindle or microtubules splaying from mid-spindle or pole regions) in metaphase-arrested TBB-1-only spindles in the *GFP::TBA-2; mCherry::H2B* genetic background (Supplemental Figure 7C-F). As such abnormalities were not detected in immunofluorescence experiments using untagged strains, we infer that GFP::TBA-2 may contribute to defects revealed in the context of meiotic arrest.

## Discussion

The research reported here was initiated with the goal of understanding how different β-tubulin isotypes contribute to assembly and function of oocyte meiotic spindles, which form and segregate chromosomes in the absence of centrosomes. Collectively, our data support a model in which β-tubulin isotype composition helps to maintain a balance between microtubule crosslinking and severing activities during oocyte meiosis in order to reliably establish spindle bipolarity, maintain spindle structural integrity during stress, and achieve normal dynamics of chromosome separation (Figure 7).

**Figure 7.**
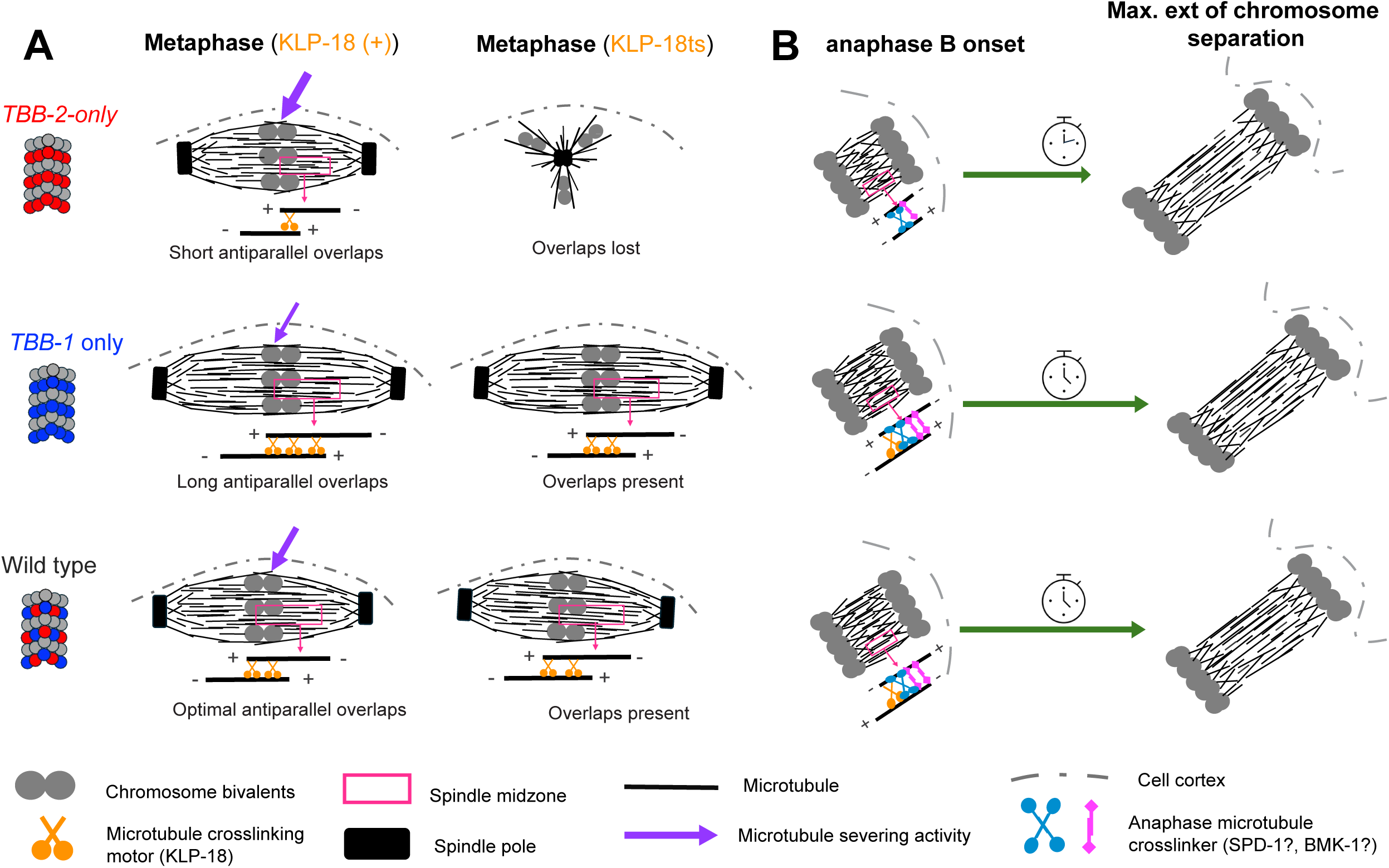
Model depicting inferred contributions of β-tubulin isotype composition to bipolar spindle assembly and anaphase chromosome separation. (**A**) We hypothesize that assembly of bipolar oocyte spindles requires a balance between microtubule crosslinking and severing activities, and that β-tubulin isotype composition helps to maintain this balance. When only the TBB-2 isotype is present, spindles are shorter due to high katanin-mediated microtubule severing activity leading to shorter antiparallel microtubule overlaps in the mid-spindle. When KLP-18 microtubule crosslinking activity is compromised, TBB-2-only spindles collapse into a monopole due to insufficient antiparallel overlap. Conversely, reduced microtubule severing activity in TBB-1-only spindles results in longer antiparallel overlaps, which are sufficient to maintain spindle bipolarity when KLP-18 activity is compromised. Presence of both TBB-1 and TBB-2 isotypes in wild type oocytes results in an optimal balance between microtubule severing and crosslinking activities. (**B**) We propose that during anaphase B, shorter microtubules in TBB-2-only spindles leads to shorter antiparallel overlaps and reduced microtubule crosslinking, making these spindles less resistant to activities of sliding motors and thereby enabling rapid spindle elongation. In contrast, microtubule crosslinking is robust in both TBB-1-only and mixed-isotype spindles due to longer antiparallel overlaps, making these spindles more resistant to sliding and hence slowing spindle elongation.

Our first evidence for differential contributions of the TBB-1 and TBB-2 isotypes came from experiments in the *klp-18ts* genetic background, in which the microtubule crosslinking motor KLP-18 is compromised (Connolly et al. 2014). Our analysis revealed a spindle assembly vulnerability in worms expressing only the TBB-2 isotype that results in a high incidence of monopolar oocyte spindles, and we further showed that this monopolar spindle phenotype could be suppressed by reducing katanin susceptibility or activity.

A model depicting our interpretation of these findings is presented in Figure 7A. We propose that increased susceptibility of TBB-2-only microtubules to katanin-mediated severing leads to shorter microtubules in the assembling meiotic spindle, which in turn results in shorter regions of overlap between crosslinked antiparallel microtubules. When crosslinking is further compromised (by the *klp-18ts* mutation), there is insufficient overlap between antiparallel microtubules to support bipolar spindle organization. Several additional observations provide further support for this model. First, we observed reduced localization of katanin around chromosomes and longer spindle lengths when only the TBB-1 isotype was present; these findings are consistent with reduced severing leading to longer microtubules in mid-spindles, thereby providing an opportunity for increased antiparallel microtubule overlap. Further, analysis of the organization of *klp-18 RNAi*-induced monopolar spindles in the isotype substitution strains provided evidence for longer microtubule bundles in TBB-1-only oocytes relative to TBB-2-only oocytes.

While TBB-1-only microtubules perform better than TBB-2-only microtubules in assembling bipolar meiotic spindles when KLP-18-mediated cross-linking is compromised, our data indicate that there is nevertheless a downside to lacking the TBB-2 isoform. Specifically, we found that the longer meiotic spindles assembled from TBB-1-only microtubules exhibit an increased incidence of structural failure in the context of prolonged metaphase arrest. This suggests that increased antiparallel MT overlap, while helping to ensure spindle bipolarity, may ultimately render spindles vulnerable to increased outward sliding forces. We infer that mixed-isotype microtubules help to achieve an optimal balance between the microtubules and the proteins that mediate interactions between them (Figure 7).

Interestingly, the same balance of tubulin isotypes that supports robustness of acentrosomal spindle assembly and function during oocyte meiosis must later support reliable assembly and function of centrosome-organized mitotic spindles during the early embryonic cell divisions (Müller-Reichert et al. 2010). Honda et al., (2017) also uncovered differential contributions of β-tubulin isotypes to functional properties of mitotic spindles. By using a tagged plus-end-binding protein to investigate the dynamics of astral microtubules in one-cell embryos, they found that TBB-2-only microtubules were more stably elongating in this context, whereas TBB-1-only microtubules were more dynamic. At a first glance, these findings might initially appear at odds with our analysis of differential contributions of β-tubulin isotypes to oocyte meiosis, where TBB-2 is associated with shorter spindles/microtubule bundles and TBB-1 is associated with longer spindles/microtubule bundles. However, as discussed above, these meiotic spindle phenotypes are largely attributable to isotype differences affecting localization of and sensitivity to katanin, which is highly abundant during meiosis but is actively degraded at the oocyte-to-embryo transition (Pintard et al. 2003; Joly et al. 2020). Thus, the properties exhibited by microtubules of similar composition can differ substantially in the very different cellular environments in which meiotic and mitotic spindles form.

Our live imaging analysis has revealed additional hidden contributions of β-tubulin isotype composition to oocyte meiotic spindle function. First, we found that TBB-1-only, TBB-2-only, and wild type oocytes achieve bipolar metaphase spindle organization with similar timing, despite TBB-2 spindles being significantly shorter. Moreover, our examination of anaphase B revealed that TBB-2-only spindles elongate significantly faster and exhibit chromosome separation velocities 67% and 43% faster than control and TBB-1-only spindles, respectively. We speculate that shorter microtubules may also underlie this faster chromosome separation phenotype of TBB-2-only spindles (Figure 7B), perhaps by shorter anti-parallel overlap offering reduced resistance to sliding and/or increasing the likelihood that anaphase motors will smoothly slide microtubules apart. Consistent with this interpretation, it was recently shown that combinatorial depletion of microtubule crosslinking factors BMK-1/ kinesin-5 and SPD-1/PRC1 leads to accelerated anaphase B spindle elongation and enhanced chromosome separation velocity (Li et al. 2023). Alternatively, or in addition, TBB-2 may drive rapid spindle elongation through effects on microtubule dynamics (*e.g.* microtubule polymerization).

There are two major take-home messages from this work:

First, our studies reveal that oocyte meiosis can appropriately be viewed as a balancing act between microtubule severing activities and microtubule crosslinking activities. Our work provides evidence that severing and crosslinking activities must counterbalance each other to form bipolar spindles and segregate chromosomes in the absence of centrosomes and further indicates that β-tubulin isotype composition helps to maintain this crucial balance.

Second, our data contribute to a growing appreciation that superficially redundant tubulin isotype variants can make distinct contributions that become relevant under specific cellular conditions. This phenomenon has been previously shown in budding yeast, where α-tubulin isotype substitution strains expressing either Tub1 only or Tub3 only are both viable under normal growth conditions, but display differential sensitivity to the microtubule depolymerizing drug Benomyl (Nsamba et al. 2021)). Additionally, the two α-tubulin isotypes contribute antagonistic properties required for optimal spindle elongation and length control during mitosis (Nsamba et al. 2025). Likewise in fission yeast, substitution of the major α-tubulin Nda2 with minor isotype Atb2 does not affect vegetative growth, but Atb2-only cells display major defects in sporulation and meiosis (Chen et al. 2025). Further, β-tubulin isotype substitution has been shown to alter astral microtubule dynamics in the mitotic spindles of *C. elegans* one-cell embryos (Honda et al. 2017), Thus, rather than simply representing functionally interchangeable components, tubulin isotypes can confer distinct properties and behaviors to microtubules that ensure robust performance of microtubule-based structures under varying cellular conditions.

## Materials and Methods

### C. elegans strains

A list of *C. elegans* strains used in this study is provided in Table 1.

**Table 1.** *C. elegans* strains used.

| Strain | Genotype | Reference |
| --- | --- | --- |
| N2 | Wild type |  |
| VC167 | ***tbb-2(0)***<br>*tbb-2(gk130) III* | (Wright and Hunter, 2003) |
| VC364 | ***tbb-1(0)***<br>*tbb-1(gk207) III* | (Wright and Hunter, 2003) |
| SA1057 | ***TBB-1-only***<br>*tbb-2(tj41[tbb-2[tbb-1 substitution]]) III* | Honda et al., 2007 |
| SA1108 | ***TBB-1-only***<br>*tbb-1(tj59[tbb-1[tbb-2 substitution]]) III* | Honda et al., 2007 |
| EU1329 | ***klp-18ts***<br>*klp-18(or447ts) IV* | Connolly et al., 2014<br>[Maintained at 15°C] |
| AV1259 | *tbb-1(gk207) III; klp-18(or447ts) / tmC5 [F36H1.3(tmls1220)] IV* | This study |
| AV1294 | *tbb-2(tj41[tbb-2[tbb-1 substitution]]) III; klp-18(or447ts) / tmC5 [F36H1.3(tmls1220)] IV* | This study |
| AV1295 | *tbb-1(tj59[tbb-1[tbb-2 substitution]]) III; klp-18(or447ts) / tmC5 [F36H1.3(tmls1220)] IV* | This study |
| AV1301 | *tbb-2(gk130) III; klp-18(or447) / tmC5[F36H1.3(tmls1220)] IV* | This study |
| AV1314 | *tbb-1(tj59[tbb-1[tbb-2 substitution]]), tbb-2(me196 E439K) III; klp-18(or447ts) / tmC5 [F36H1.3(tmls1220)] IV* | This study |
| AV1340 | *mei-2(me198 A237T) I; tbb-1(tj59[tbb-1[tbb-2 substitution]]) III; klp-18(or447ts) / tmC5 [F36H1.3(tmls1220)] IV* | This study |
| AV1342 | *mei-2(me198 A237T), mei-1(wow21[ZF:GFP:mei-1]) I; zif-1(gk117) III* | This study |
| AV1348 | *tbb-2(me196 E439K) III*<br>[me196 is recreation of *tbb-2(sb26)*] | Lu et al., 2004. |
| AV1393 | *mei-2(me198, A237T) I*<br>[me198 is recreation of *mei-2(ct98)*] | Srayko et al., 2000. |
| OD57 | *unc-119(ed3) III;; Itls25[pAZ132; pie-1::GFP::tba-2; unc-119 (+)] IV; Itls37 [pAA64; pie-1/mCherry::his-58; unc-119 (+)]* | Dumont et al., 2010 |
| AV1361 | *tbb-2(tj41[tbb-2[tbb-1 substitution]]) III; Itls25[pAZ132; pie-1::GFP::tba-2; unc-119 (+)] IV; Itls37 [pAA64; pie-1/mCherry::his-58; unc-119 (+)]* | This study |
| AV1362 | *tbb-1(tj59[tbb-1[tbb-2 substitution]]) III; Itls25[pAZ132; pie-1::GFP::tba-2; unc-119 (+)] IV; Itls37 [pAA64; pie-1/mCherry::his-58; unc-119 (+)]* | This study |

Except where noted, worms were cultured under standard conditions (Stiernagle 2006) at 20°C.

For experiments involving *tbb-1(0); klp-18ts* or *TBB-2-only; klp-18ts* worms, homozygous mutant worms were derived from strains carrying *klp-18ts in trans* to the *tmC5* balancer.

### RNAi feeding

RNAi was done as previously described (Wolff et al., 2022). Briefly, individual RNAi clones from an RNAi feeding library (Kamath et al., 2003) were streaked on LB plates containing ampicillin and grown overnight at 37°C. The next day, single colonies were grown overnight at 37°C in LB with 100 μg/mL ampicillin for 16-18 hours. Overnight cultures were spun down, resuspended, and plated on nematode growth medium (NGM) plates containing 100 μg/mL ampicillin and 1 mM IPTG. Plates were dried in a dark space overnight at 23°C. On the same day, worms were synchronized by bleaching of gravid adults, collecting their embryos, and plating embryos on unseeded plates to hatch overnight at 20°C. The next day, newly hatched L1 worms were transferred to appropriate RNAi plates and grown to adulthood at 20°C for 4 days.

### Immunofluorescence and antibodies

Immunofluorescence was performed as described in Wolff et al., 2022. Adult gravid worms were picked into a 5 μl drop of sterile M9 (22 mM KH2PO4,42mMNa2HPO4, 85.5 mM NaCl, 1 mM MgSO4) on glass coverslips (22 x 22 mm) and were dissected to release embryos. A poly-L-lysine slide was laid on top to pick up the coverslip and then plunged into liquid nitrogen for 5 minutes. The coverslip was rapidly flicked off via razor blade and slides were fixed in cold methanol for 35 minutes at -20°C. Slides were rehydrated twice in 1X PBS with 0.1% Triton-X-100 (PBS-T) and then blocked in AbDil buffer (1X PBS with 4% BSA, 0.1% Triton-X-100, 0.02% Sodium Azide) at room temperature for at least 60 minutes. Primary antibodies were diluted in AbDil, and slides were incubated in a humid chamber overnight at 4°C. The following day, slides were washed three times with PBS-T and incubated in secondary antibodies diluted in AbDil for two hours (in the humid chamber) at room temperature. Samples were rinsed again three times in PBS-T as before and incubated in Hoechst (D1306 Invitrogen at 1:1000 in PBS-T) for 12-15 minutes at room temperature. The samples were washed one final time twice in PBS-T and mounted in 5-6 μl of mounting media (0.5% p-phenylenediamine, 20 mM Tris-Cl, pH 8.8, 90% glycerol), a coverslip (22 x 22 mm #1) was gently laid on top and then sealed with nail polish. Mounted slides were stored at 4°C.

Primary antibodies used in this study were rabbit-α-ASPM-1 (1:5000, [Wignall and Villeneuve 2009]), rabbit-α-MEI-1 (1:100, gift from Frank McNally), rabbit-α-MEI-2 (1:100, [(McNally et al. 2014)]), and directly conjugated mouse-α-Tubulin-FITC (1:500, DM1α, Sigma). Alexa-fluor conjugated secondary antibodies (Invitrogen) were used at 1:500. Fixed imaging was performed at 4°C on an Inverted Zeiss LSM 880 Laser Scanning Confocal Microscope with a 63x (Plan APO for AiryScan) oil (NA = 1.58) objective. The microscope is equipped with a Piezo-driven motorized stage (PZ-2000 XYZ, Applied Scientific Instrumentation). Z-stacks were obtained at 0.3-μm increments. Undeconvolved images were processed with ImageJ. Unless otherwise stated, images in the paper are shown as maximum intensity projections. The LSM 880 microscope is housed in the Cell Sciences Imaging Facility located in and supported by the Beckman Center for Molecular and Genetic Medicine.

### Embryonic viability assays

Except where noted, embryonic viability, X-chromosome missegregation and brood sizes were measured at 20°C for 12 individual worms of each genotype. Individual late L4 hermaphrodites were singled onto seeded NGM plates containing a small bacterial lawn. Individual worms were transferred daily (2x) to a fresh plate. Plates were scored 24 hours after removal of the parent for the number of eggs and hatched larva; plates were scored again 3 days later to assess numbers of adult hermaphrodites and males.

### CRISPR generation of *tbb-2(E439K)* and *mei-2(A237T)* alleles

CRISPR editing was performed as described (Paix et al. 2015). sgRNA and PAM sites for all targets were selected using the CRISPOR website https://crispor.gi.ucsc.edu/. A list of crRNAs and primers used are provided in Table S4. An injection mixture of target gene crRNA and repair oligo, co-CRISPR marker *dpy-10* repair oligo (Arribere et al., 2014, IDT), *dpy-10* crRNA (GCTACCATAGGCACCACGAG, IDT), trRNA (IDT) and Cas9-NLS nucleases (IDT) was injected into young adults of the appropriate genotypes. Appropriate edits were identified by PCR screening and confirmed by Sanger sequencing.

### Live *in utero* imaging

L4 larvae were incubated at 20°C overnight on seeded NGM plates, and 15-20 young adult hermaphrodites were picked into 1.5 μl of a worm soaking solution containing 20 mM Serotonin (H7752-1G,Sigma-Aldrich), 83mM tetramisole hydrochloride (Levamisole, Sigma L9756) and 10% tricaine (prepared in water and kept on ice) on the bottom of #1.5 Ibidi 35 mm dish (50305805 Fisher Scientific). 0.75 μl of 0.1 μm polystyrene microspheres (08691 Polysciences) was added to the mixture and worms were left to immobilize for 10 minutes. A thin agarose pad (5% agarose in water and soaked in 40 μl of soaking solution briefly after preparation) was gently dried with a Kimwipe and then inverted and laid on top of the worms using tweezers to create a dish-worm-pad sandwich. A wet Kimwipe was rolled around the bottom of the Ibidi dish and the lid placed on top to create a humid chamber. The dish was placed on the top of an inverted spinning disk microscope and imaged from below. Meiotic oocytes before onset of ovulation were identified by bright field microscopy before initiating time-lapse fluorescence imaging.

Fluorescence imaging was performed at 20°C on a Nikon Ti-E Inverted spinning disk confocal microscope (Nikon Instruments) using a 60x PLAN APO oil objective (NA = 1.4) controlled using NIS Elements software (Nikon). Movies of oocytes/embryos expressing GFP::TBA-2 and mCherry::H2B were captured using an Andor DU-897 X-6910 camera controlled using NIS Elements software (Nikon). Z-series time-lapse images (Z-stack containing the entire oocyte, 2-μm step size; Laband et al., 2018) were collected at 12 s intervals. The 488 nm and 561 nm channels were imaged simultaneously at 5% and 10% power, respectively, at 250 ms exposure time. Focus was adjusted manually during time-lapse imaging.

### Analysis of meiosis I timing

To assess progression through meiosis I, we designated the appearance of microtubules around chromosomes following nuclear envelope breakdown to mark onset of meiotic spindle assembly (t=0) and collected z-stacks every 12 seconds. Cell cycle stages and landmark events (illustrated in Figure 6A) were defined as previously described by pioneering live-imaging studies of *C. elegans* oocyte meiosis (Chuang et al. 2020). Metaphase is when chromosomes are fully congressed on bipolar spindles; Anaphase A onset is when the spindle has shortened to its shortest length and bivalents have separated into homologs and begun moving toward opposite poles; Anaphase B onset is when the chromosome masses have reached the outside of the spindle microtubules and the spindle starts to elongate; Maximum extent of chromosome separation is when the spindle has reached maximum length and chromosome masses are separated by maximum distance, before actinomyosin ring contraction causes the spindle to bend away from the cortex; End of meiosis I is defined by polar body extrusion.

Chromosome separation over time and Anaphase B chromosome separation velocity were analyzed as previously described (Li et al. 2023). For each spindle, distances between separating homologs (anaphase I) were measured in 3D in ImageJ by drawing a line connecting the centers of both chromosome masses and measuring the length of the line in μm; in cases where the spindle became bent and GFP::TBA-2-labelled spindle microtubules were clearly visible, a segmented line was drawn along the bent spindle to connect the two chromosome masses. Measurements were made every 24 seconds (2 frames) for a period of 200-250 seconds, until maximum extent of chromosome separation was achieved. We note that TBB-2-only spindles achieved maximum extent of chromosome separation in under 200 seconds. To represent anaphase chromosome separation over time, chromosome separation distances (every 24 seconds since anaphase A onset (t=0)) for multiple spindles were averaged together to generate the cumulative line profiles shown in Fig. 6E. Anaphase B chromosome separation velocities for each spindle (for the first 150 seconds in Fig 6F and for the entire anaphase B duration Fig S6D) were calculated as follows: the distance between chromosome masses at t=0 (anaphase B onset) was subtracted from distance between chromosome masses at the end point, and this value was divided by the elapsed time (in seconds) to obtain separation velocity in μm per second.

#### GFP::TBA-2 and mCherry::H2B fixed fluorescence imaging

Worms were prepared, dissected and fixed as for immunofluorescence experiments. After 35 minutes in methanol, slides were washed twice in 1X PBST, mounted in mounting media (0.5% p-phenylenediamine, 20 mM Tris-Cl, pH 8.8, 90% glycerol), a coverslip (22 x 22 mm #1) was gently laid on top and then sealed with nail polish. Samples were imaged on a Zeiss LSM 880 Confocal microscope with a 63X oil (NA = 1.58) objective.

### MEI-1/2 Signal Distribution Analysis

Analysis was performed in ImageJ using Sum projected images of spindles oriented roughly perpendicular to the Z axis. For each spindle, a “line” wide enough to capture the full width of the spindle was drawn along the long axis of the spindle, connecting the poles; pixel values at each position were extracted in ImageJ and recorded in Excel. Identical lines were drawn in an area of the cytoplasm adjacent to the spindle to extract background signal values, which were averaged to obtain mean background signal. For each position along the spindle, background-subtracted signal values were calculated, and signals were further internally normalized by dividing by the mean background-subtracted signal for the entire spindle. Finally, the normalized signal distribution for each spindle was then plotted against normalized spindle length (obtained by diving distances along the spindle by the maximum spindle length) in Excel to generate line profiles in Figs 4C and S4C.

To compare distributions between genotypes, we generated an additional metric for each spindle, *i.e.* the average normalized signal for the central spindle (from 0.4 to 0.6 along the normalized spindle length).

### Statistical methods

Unless otherwise stated, two-tailed Mann-Whitney P values were calculated in GraphPad Prism 10. Fisher’s exact tests were used to analyze contingency tables for categorical data using https://www.graphpad.com/quickcalcs/.

### Quantification of other spindle features

#### Spindle length

Analyzed in 3D using ImageJ. For fixed immunofluorescence images in Fig 5B, the center of each pole was defined using the XYZ coordinates of the ASPM-1 signal, as it represents the point where the dense pole-associated microtubules terminate at the ends of the spindle. For the live images in Fig 6B and S7B, the ends of spindles were estimated from GFP::TBA-2 fluorescence, with poles defined as the point where bright GFP::TBA-2 fluorescence terminates. Spindle lengths (in μm) were determined from the XYZ coordinates using the formula:

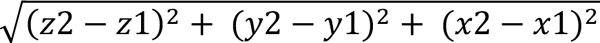

#### ASPM-1 and KLP-18 signal measurements

Image analysis was performed in ImageJ using sum-projected images of meiosis I spindles acquired identically at similar laser power. For each spindle, a sum projection of z-sections encompassing the full spindle was generated, an elliptical region of interest (ROI) was drawn around the pole in the channel of the sum projection, and the integrated density (mean pixel intensity x area) was recorded for the channel of interest. To calculate the background for each channel of interest, the same ROI was shifted to an area of the cytoplasm next to each pole and integrated density measurements were recorded. For each spindle, the background measurements were then subtracted from pole measurements, and the two background-subtracted poles were averaged together to obtain total signal. Each data point represents one spindle.

For KLP-18 normalized signal, the background-subtracted KLP-18 signal was divided by the background-subtracted tubulin signal to normalize for variability in staining.

#### Microtubule bundle length

Distances (in μm) from the base of microtubule bundle to mid-bivalent were measured in 3D using XYZ coordinates as for spindle length measurements. The base of the bundle is defined as a region of high microtubule density that colocalizes with high ASPM-1 signal in the area of the monopole; groups of microtubule bundles emanating from the base form lateral associations with a given bivalent. For each monopolar spindle, a line was drawn connecting the base to mid-bivalent for each bivalent, and all measurements were averaged together to obtain an average distance for a given spindle.

#### Midspindle Angle

Midspindle angles were measured using maximum intensity projections of spindles in which both poles were visible. The center of the spindle was identified as the area containing mCherry::H2B fluorescence corresponding to the congressed chromosomes. Midspindle angle was measured in ImageJ by drawing a line connecting the center of spindle to the center of each spindle pole. The resulting data were binned (5^°^ increments) and visualized in R studio

## Supporting information

Nsamba 2025 SI combined

## Data availability

All data and materials used in this study are available upon request from A.M. Villeneuve.

## Acknowledgments

We are grateful to Chantal Akerib for expert advice regarding genetic analyses and assistance with strain constructions. We thank members of the Jessica Feldman lab, in particular Jeremy Magescas and Wenzhe Li for expert advice, technical assistance and support for live *in utero* imaging and for providing access to their microscope. We thank Ivana Cavka, Kitty Lee, and members of the Villeneuve lab for technical assistance and discussions. We thank Asako Sugimoto, Sarah Wignall, Francis McNally, and the Caenorhabditis Genetics Center (funded by NIH Office of Research Infrastructure Programs P40 OD010440) for reagents and strains. This work was supported by the Stanford University Cell Sciences Imaging Core Facility (RRID:SCR_017787) and by National Institutes of Health grant R35GM126964 to A.M.V.

## Author Contributions

E.T. Nsamba: Conceptualization, Investigation, Methodology, Formal analysis, Validation, Writing (original draft), Writing ( review and editing). A.M. Villeneuve: Conceptualization, Formal Analysis, Methodology, Validation, Writing (original draft), Writing ( review and editing), Supervision, Funding acquisition.

