## Supplementary material for "Differential contributions of β-tubulin isotypes to acentrosomal oocyte meiosis in *C. elegans*": Nsamba 2025 SI combined

**Includes;**

Supplemental Figure Legends

Supplemental Tables

Supplemental Figures S1- S7

**Supplemental Figure Legends:**

Figure S1. **Loss of TBB-1 exacerbates meiotic spindle assembly defects caused by *klp-18ts*.** Representative IF images of disorganized metaphase I arrested spindles observed in the *TBB-2-only; klp-18ts* mutant background. In contrast to monopolar or bipolar spindles, which have one or two poles marked by ASPM-1 foci, respectively, spindles classified as "disorganized" may exhibit no ASPM-1 foci (apolar), more than two ASPM-1 foci (multipolar), or have ASPM-1 distributed broadly within the spindle. Scale bars, 2.0 µm.

Figure S2. **Analysis of spindle morphologies in unsynchronized embryos harboring *klp-18ts* mutation. (A)** Representative images of GFP and mCherry fluorescence in unsynchronized metaphase/metaphase-like spindles in fixed *klp-8ts* single mutant and *TBB-2-only; klp-8ts* double mutant embryos expressing GFP::TBA-2 and mCherry::H2B. Monopolar or disorganized spindles were classified as "metaphase-like" if bivalents were unseparated and were flanked by discernable microtubule bundles. **(B)** Quantification of spindle morphologies observed in A. Similar to data from immunofluorescence images of metaphase I arrested spindles, loss of TBB-1 exacerbates spindle bipolarity defects in the *klp-18ts* mutant background. Analysis of spindle bipolarity as in Figure 1G, $p=0.0001$. **(C)** Representative GFP::TBA-2 and mCherry::H2B fluorescence images showing anaphase-like spindle morphologies in fixed *klp-8ts* single mutant and *TBB-2-only; klp-8ts* double mutant embryos. Anaphases were scored as normal based on the presence of two separated chromosome masses of comparable size flanking a dense microtubule array. Spindles were classified as "abnormal anaphase" if they contained >2 chromosome masses of distinct sizes at the periphery of a dense microtubule array, a single chromosome mass adjacent to a compact dense microtubule array, or a disorganized set of chromosome masses within a condensed spindle lacking a central dense microtubule array. **(D)** Quantification of anaphase spindle morphologies in C. *TBB-2-only; klp-8ts* embryos exhibit defects in anaphase spindle assembly and aberrant chromosome segregation likely arising from earlier defects in spindle bipolarity. In B and C, both meiosis I and meiosis II spindles were grouped together. Scale bars in all images = 2.0 µm

Figure S3. **No evidence for a difference in KLP-18 localization between TBB-1-only and TBB-2-only oocyte meiotic spindles**. **(A)** Representative IF images of metaphase I spindles showing variability of KLP-18 IF signals in the embryos examined. Scale bar = 2.0 µm. **(B)** Quantification of KLP-18 fluorescence IF signals normalized to tubulin IF signals. Horizontal lines indicate median. n = total number of embryos analyzed per genotype. ns, not significant; $*p < 0.05$.

Figure S4. **TBB-1-only spindles exhibit reduced localization of katanin around chromosomes relative to the poles. (A)** MEI-1 immunofluorescence in metaphase I-arrested oocytes showing differential chromosome association. Shown are DNA (blue), tubulin (green), MEI-1 (red). Scale bar = 2.0 μm. **(B)** Analysis schematic for measuring MEI-1 distribution across spindle axis. Positions 0.4 – 0.6 represents central region in the vicinity of chromosomes. **(C)** Individual line traces of normalized MEI-1signal intensity from measurements in B from spindles of the indicated genotypes. TBB-1-only spindles exhibit reduced MEI-2 central spindle localization. **(D)** Quantification of central spindle MEI-1 signal (positions 0.4–0.6); each data point represents an average of internally normalized values per spindle. Horizontal lines indicate median. Data were analyzed using two-tailed Mann-Whitney test; ns, not significant; ***p < 0.001; ****p < 0.0001.

Figure S5. **ASPM-1 localization and spindle pole organization are largely normal in TBB-1-only and TBB-2-only spindles. (A)** Representative IF images of metaphase I arrested spindles showing ASPM-1 localization and pole width differences for the indicated genotypes; scale bars = 2.0 μm. **(B)** Quantification of ASPM-1 pole-associated signal shown in A. **(C)** Quantification of pole widths shown in A; schematic depicts how pole widths were measured. Although both TBB-1-only and TBB-2only spindles had slightly narrower poles than wild-type, these modest differences contrasted sharply with the more pronounced defects detected in the *gfp::mei-1; mei-2(A237T) and tbb-2(E439K) mutants. For B and C,* error bars indicate standard error of the mean. n = number of embryos analyzed. Data were analyzed using two-tailed Mann-Whitney test; ns, not significant; *p < 0.05; ***p < 0.001; ****p < 0.0001.

Figure S6. **Additional analysis for live imaging presented in Figure 6. (A-C)** As in Figure 6C, with additional landmark-to-landmark intervals included. **(D)** As in Figure 6F, with a different endpoint.

Figure S7. **TBB-1-only spindles exhibit increased susceptibility to structural failure during prolonged arrest. (A)** Representative fixed images of metaphase I arrested spindles in control, TBB-2-only and TBB-1-only oocytes expressing GFP::TBA-2 and mCherry::H2B. **(B)** Quantification of meiotic spindle length in A; TBB-1-only spindles are significantly longer than wild type and TBB-2 only spindles. Error bars indicate standard deviation. **(C)** GFP::TBA-2 and mCherry::H2B fixed fluorescence images illustrating bent midspindle morphology. **(D)** Quantification of spindle angle as illustrated in C. TBB-1-only spindles are significantly bent (p<0.0001 vs control, and p<0.0001 vs TBB-2-only; two-tailed Mann-Whitney test), reflecting structural failure. **(E)** GFP::TBA-2 and mCherry::H2B fixed fluorescence images illustrating splaying of microtubules from mid-spindle and pole regions. **(F)** Quantification of microtubule splaying phenotype illustrated in E. Scale bars in all images = 2.0 μm.

**Table S1**. Progeny production by worms carrying *klp-18(or447ts)* and/or β-tubulin mutations.

| **Maternal genotype** | **% hatching (20°) (no. of embryos)** | **% viability (20°) (no. of adults)** | **% Males (no. of adults)** | **Average no. of eggs laid (no. of worms)** |
|---|---|---|---|---|
| *tbb-1(0)* | 98.5 (2526) | 96.7 (2440) | 0.1 (2440) | 253 (10) |
| *tbb-2(0)* | 60.0 (1908) | 52.1 (995) | 2.0 (995) | 127 (15) |
| *klp-18ts* | 53.4 (1959) | 45.0 (881) | 7.4 (881) | 163 (12) |
| *tbb-1(0); klp-18ts* | 2.1 (2091) | 0.5 (10) | 50 (10) | 209 (10) |
| *tbb-2(0); klp-18ts* | 26.0 (1672) | 13 (215) | 7.0 (215) | 139 (12) |
| *TBB-1-only; klp-18ts* | 60.3 (1997) | 51.3 (1025) | 5 (1025) | 166 (12) |
| *TBB-2-only; klp-18ts* | 3.2 (1766) | 1.2 (21) | 24 (21) | 147 (12) |

**Table S2**. *tbb-2(E439K)* suppresses meiotic defects of *TBB-2-only; klp-18(or447ts)*

| **Maternal genotype** | **% hatching (no. of embryos)** | | **% viability (no. of adults)** | | **% Males (no. of adults)** | | **Average no. of eggs laid (no. of worms)** | |
|---|---|---|---|---|---|---|---|---|
| | **20°** | **25°** | **20°** | **25°** | **20°** | **25°** | **20°** | **25°** |
| *klp-18ts* | 75.6 (1744) | 0.8 (360) | 73.3 (1279) | 0.3 (1) | 2.3 (1279) | 100 (1) | 159 (11) | 60 (6) |
| *TBB-2-only; klp-18ts* | 3.6 (2341) | 0.9 (774) | 1.8 (42) | 0.1 (1) | 21 (42) | 0 (1) | 195 (12) | 129 (6) |
| *tbb-2(E439K), TBB-2-only; klp-18ts* | 82.7 (2001) | 11.9 (494) | 78.2 (1564) | 9 (44) | 1.4 (1564) | 2.3 (44) | 167 (12) | 82 (6) |

**Table S3**. Suppression of meiotic defects of *TBB-2-only; klp-18(or447ts)* by *mei-2(A237T)* mutation causing reduced katanin activity.

| **Maternal genotype** | **% hatching (20°) (no. of embryos)** | **% viability (20°) (no. of adults)** | **% Males (no. of adults)** | **Average no. of eggs laid (no. of worms)** |
|---|---|---|---|---|
| *mei-2(A237T)* | 99.1 (3443) | 99.1 (3413) | 0.0 (3413) | 287 (12) |
| *klp-18ts* | 82.6 (1965) | 78.2 (1537) | 2.3 (1537) | 164 (12) |
| *TBB-2-only; klp-18ts* | 4.0 (2190) | 1 (21) | 19 (21) | 183 (12) |
| *mei-2(A237T); TBB-2-only; klp-18ts* | 74.8 (2304) | 66.1 (1523) | 0.7 (1523) | 192 (12) |

**Table S4:** Reagents for CRISPR-Cas9 editing

| Allele | Target gene | Sequence description | Sequence |
|---|---|---|---|
| *me196 (E439K)* | *tbb-2* | sgRNA | actgccgaagacgacgtcga |
| | | Repair template | CAACAATACCAAGAAGCAACTGCCGAAGACGACGTCGACGGATACGCTAAGGGAGAAGCTGGAGAGACTTACGAATCGGAGCAA |
| | | Forward primer | CCTTCGTTGGAAACTCGACC |
| | | Reverse primer | TCAAGCGTATCTCAGAGCAG |
| *me198 (A237T)* | *mei-2* | sgRNA | tcggcgaattggagtcgatg |
| | | Repair template | GATTTGCATCCACCAAAACTCGGCGAATTGGAGTCGACGTTGTTGCTGAAGAGAGAGCTGCTAAAACAACAGAGTGCATTCACAATTTTCGCAAAATAGTT |
| | | Forward primer | CTTGGTGTCTCAACTCGTGC |
| | | Reverse primer | AATCAGTTCCTCCACGAGG |

### Examples of disorganized spindles

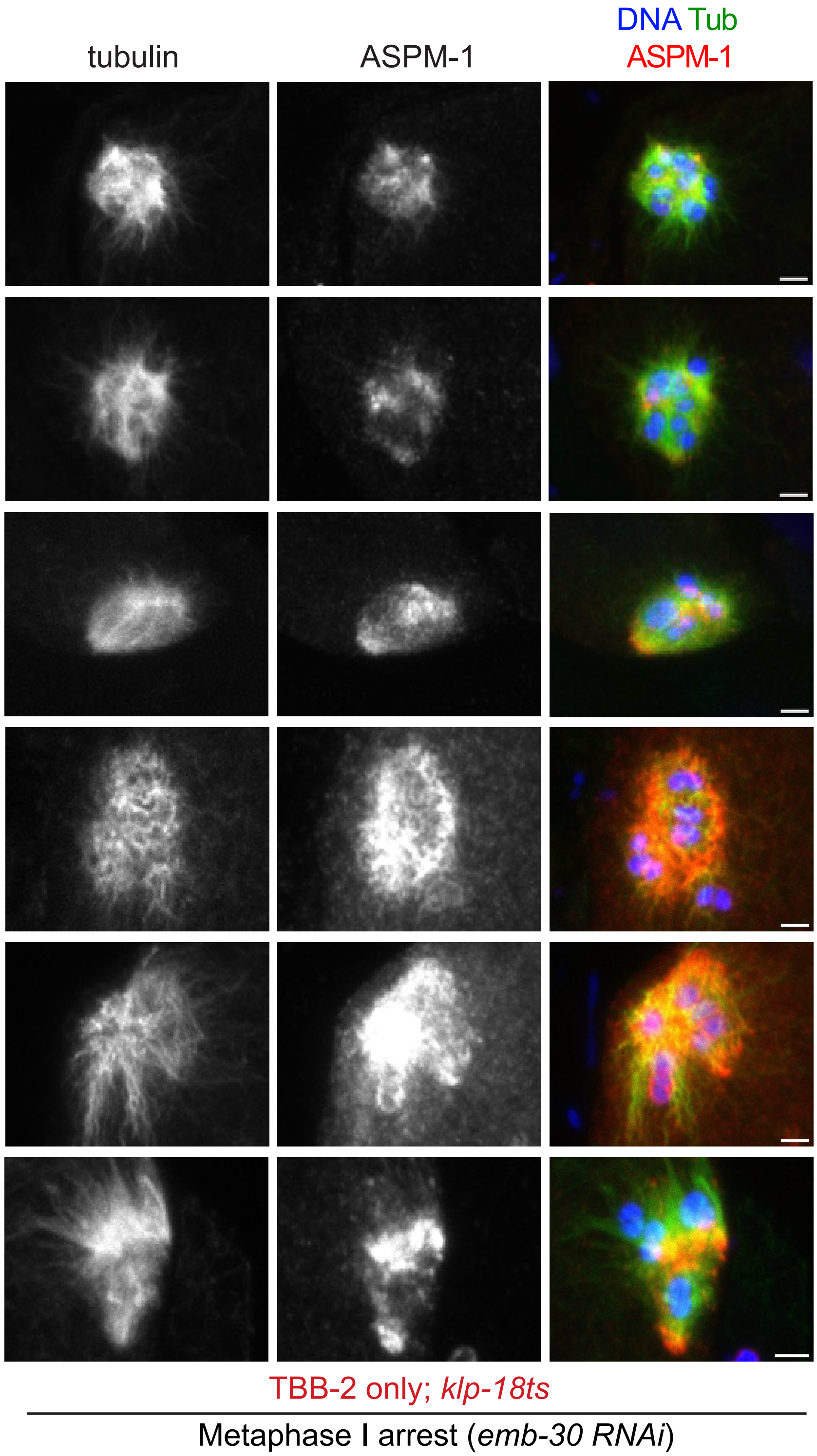

Supp. Figure 2

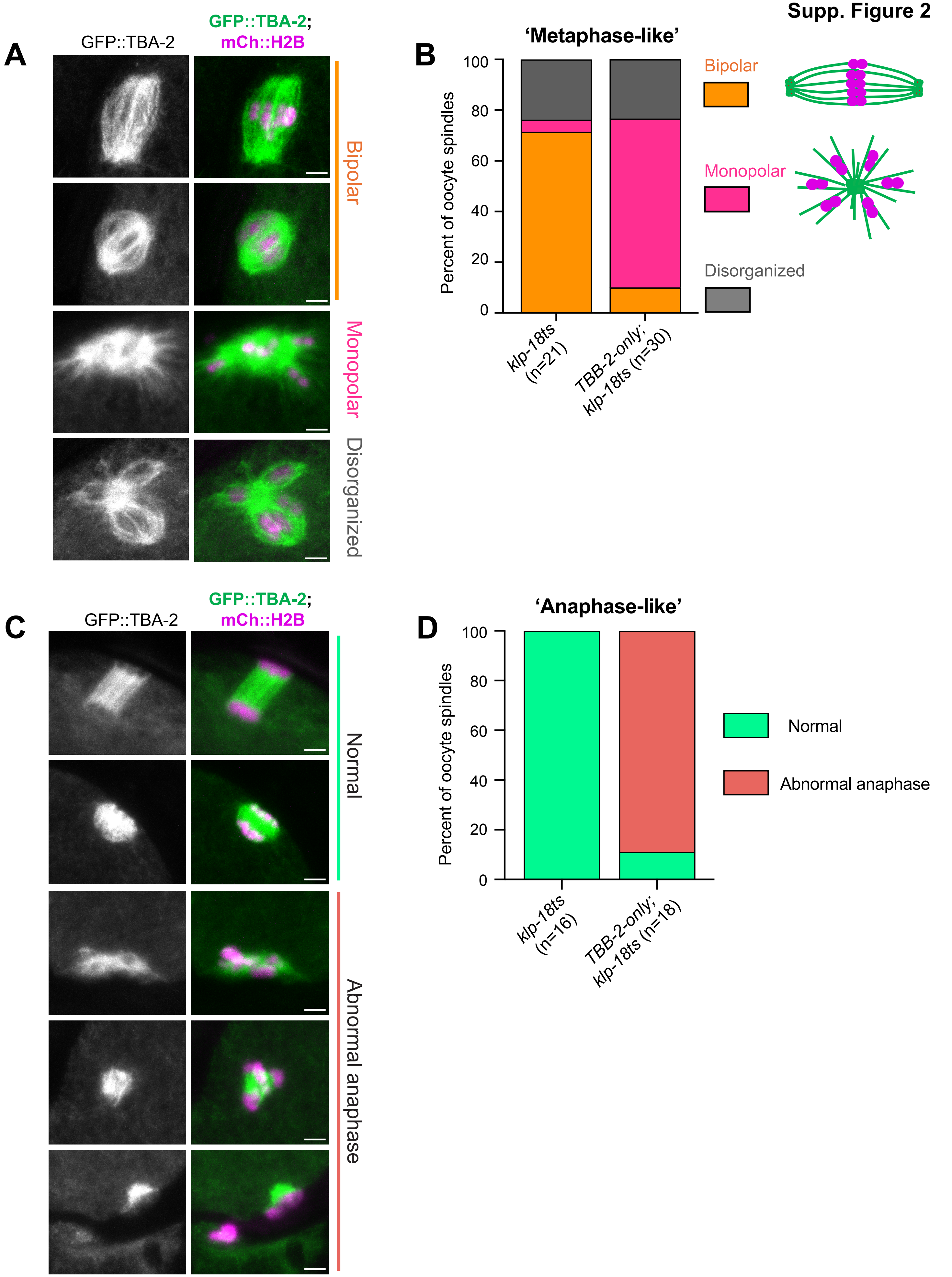

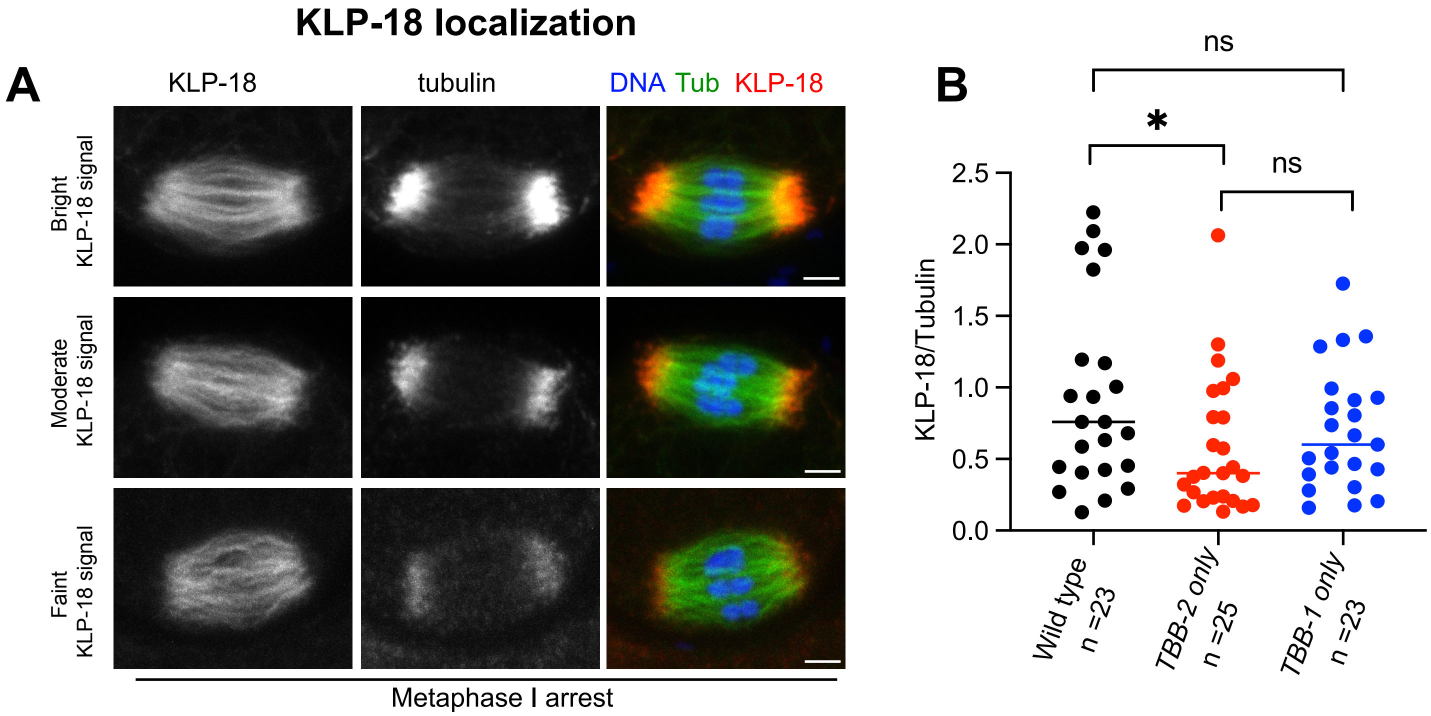
KLP-18 localization
A
KLP-18
tubulin
DNA Tub KLP-18
Bright KLP-18 signal
Moderate KLP-18 signal
Faint KLP-18 signal
Metaphase I arrest
B
ns
*
ns
KLP-18/Tubulin
2.5
2.0
1.5
1.0
0.5
0.0
Wild type n =23
TBB-2 only n =25
TBB-1 only n =23

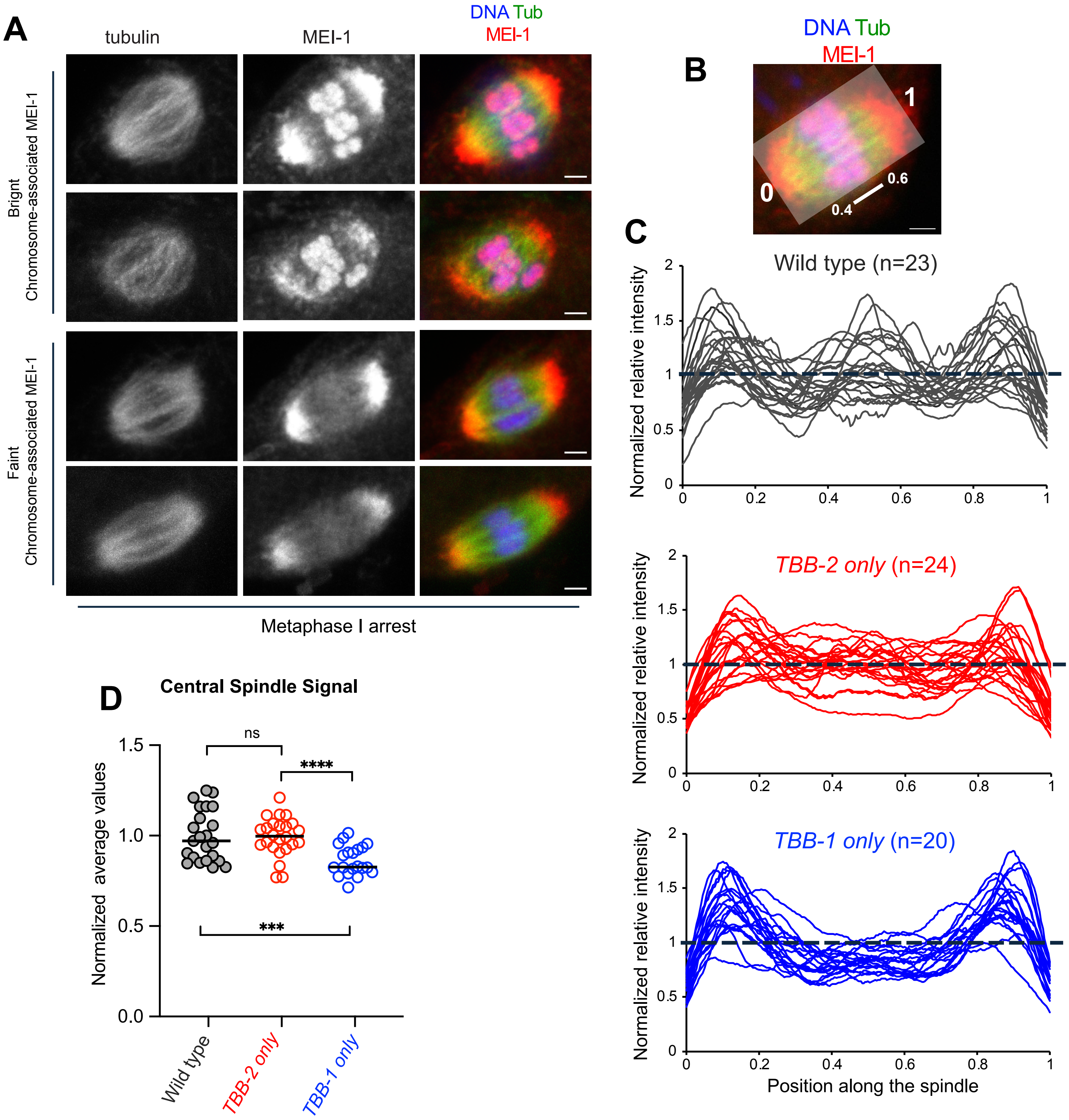
A
tubulin
MEI-1
DNA Tub
MEI-1
Brignt
Chromosome-associated MEI-1
Faint
Chromosome-associated MEI-1
Metaphase I arrest
B
DNA Tub
MEI-1
1
0
0.6
0.4
C
Wild type (n=23)
TBB-2 only (n=24)
TBB-1 only (n=20)
Normalized relative intensity
Position along the spindle
D
Central Spindle Signal
Normalized average values
ns
****
***
Wild type
TBB-2 only
TBB-1 only

**Supp. Figure 5**

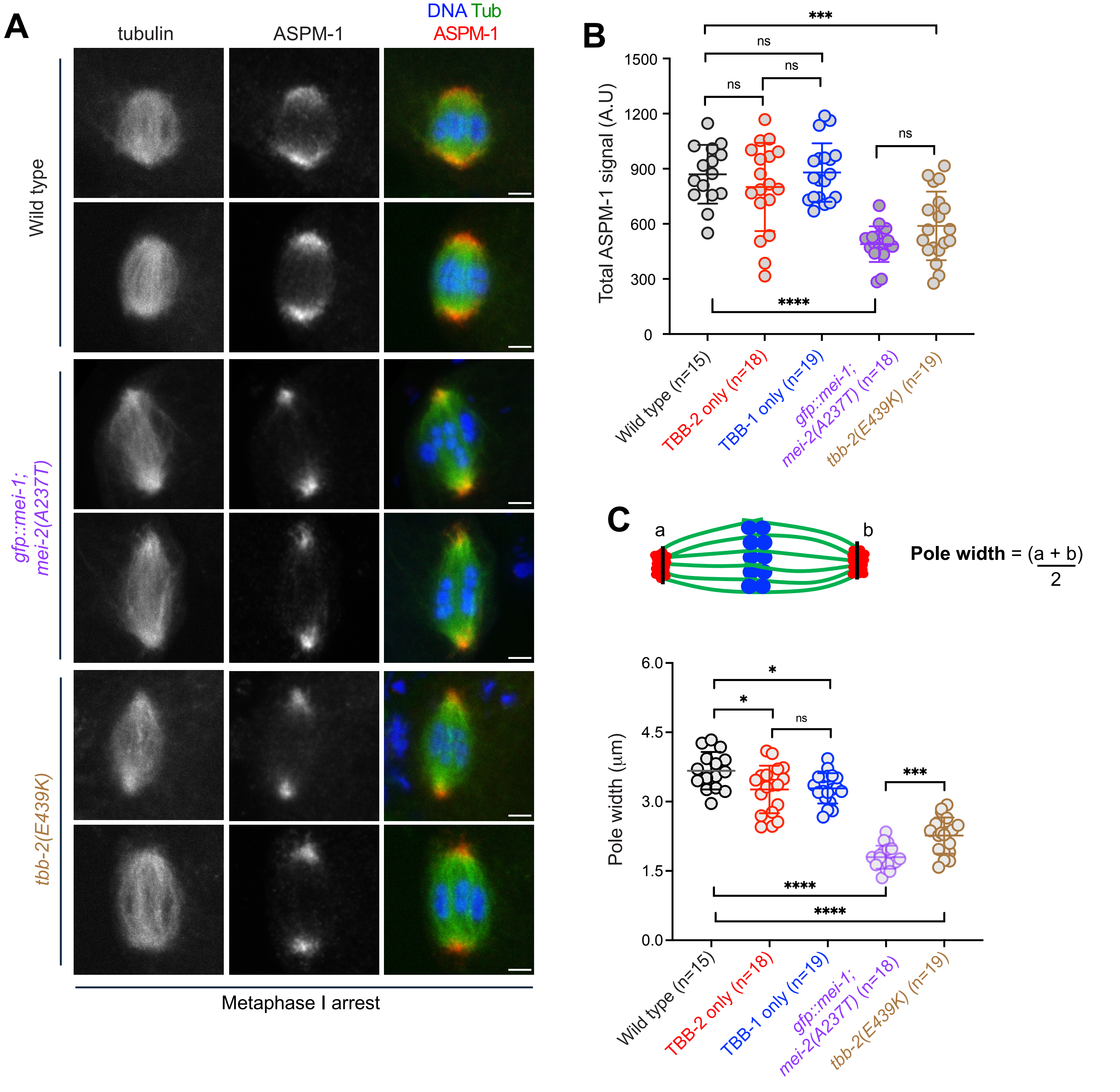

**Supp. Figure 6**

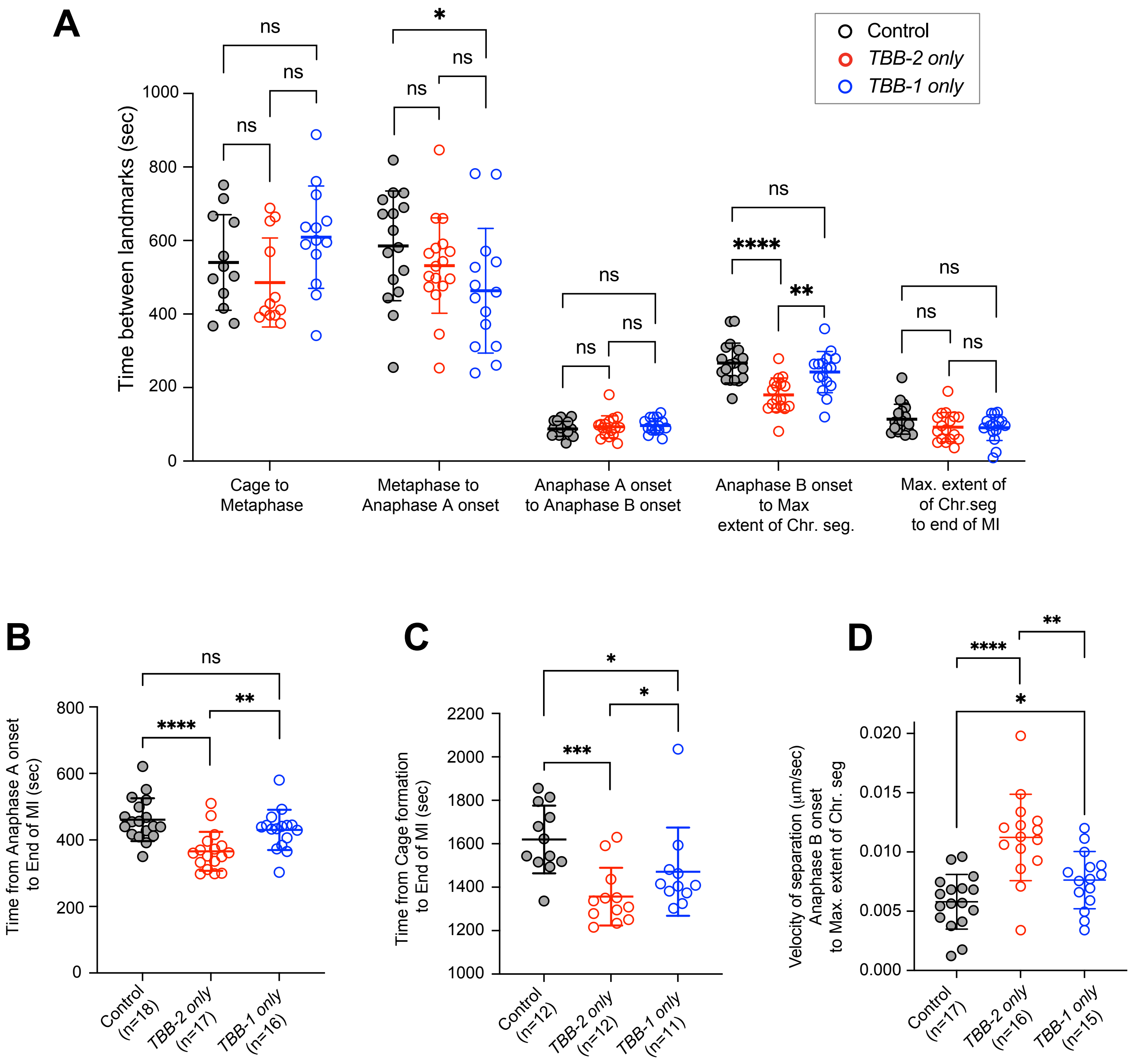

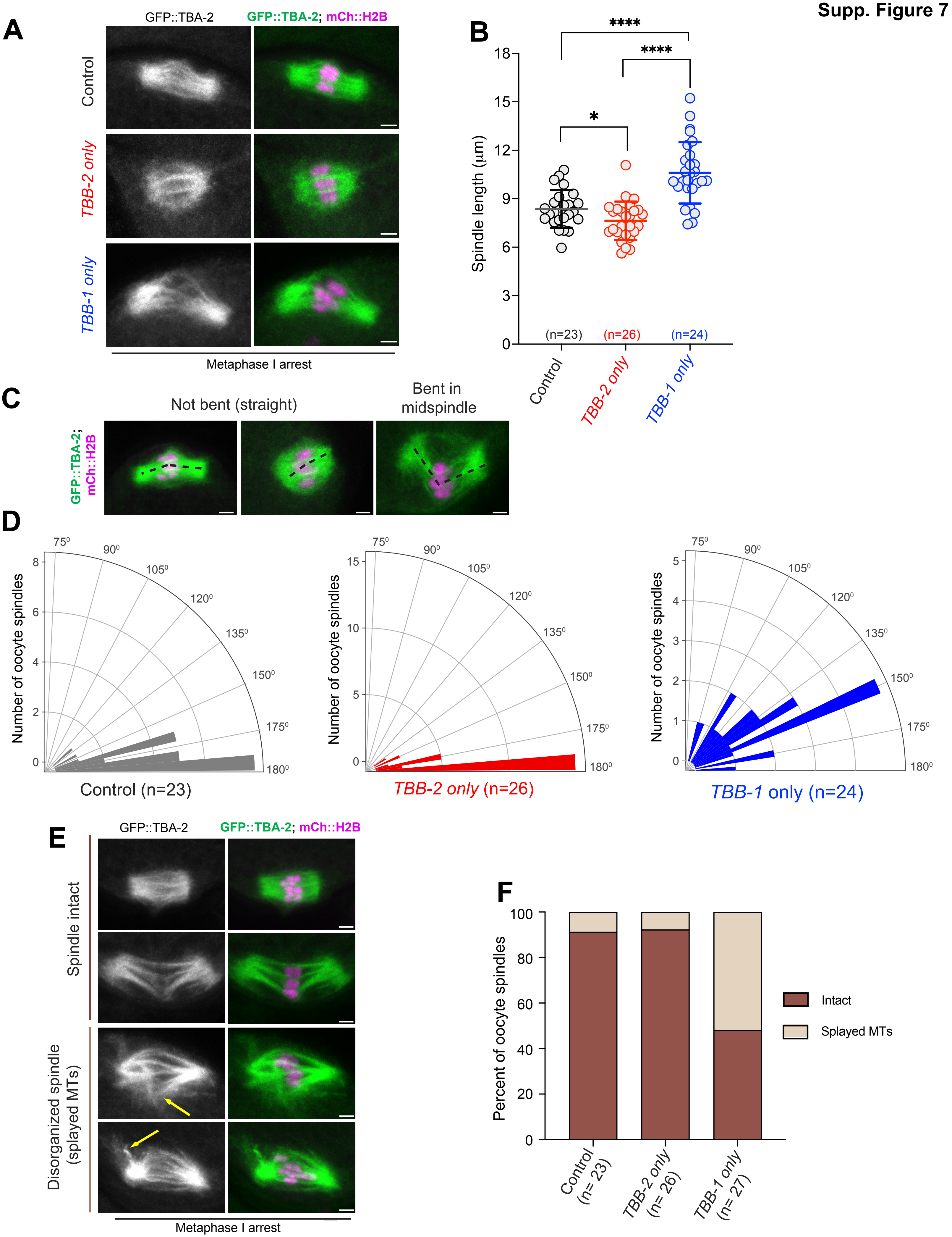
A
GFP::TBA-2
GFP::TBA-2; mCh::H2B
Control
TBB-2 only
TBB-1 only
Metaphase I arrest
B
****
****
*
Spindle length (µm)
18
15
12
9
6
3
0
(n=23)
(n=26)
(n=24)
Control
TBB-2 only
TBB-1 only
C
Not bent (straight)
Bent in midspindle
GFP::TBA-2; mCh::H2B
D
Number of oocyte spindles
Control (n=23)
TBB-2 only (n=26)
TBB-1 only (n=24)
E
GFP::TBA-2
GFP::TBA-2; mCh::H2B
Spindle intact
Disorganized spindle (splayed MTs)
Metaphase I arrest
F
Percent of oocyte spindles
100
80
60
40
20
0
Control (n= 23)
TBB-2 only (n= 26)
TBB-1 only (n= 27)
Intact
Splayed MTs
